# Structured memory representations develop at multiple time scales in hippocampal-cortical networks

**DOI:** 10.1101/2023.04.06.535935

**Authors:** Arielle Tambini, Jacob Miller, Luke Ehlert, Anastasia Kiyonaga, Mark D’Esposito

**Affiliations:** Center for Biomedical Imaging and Neuromodulation, Nathan Kline Institute for Psychiatric Research, Orangeburg, NY; Department of Psychiatry, New York University Grossman School of Medicine, New York, NY; Wu Tsai Institute, Department of Psychiatry, Yale University, New Haven, CT; Department of Neurobiology and Behavior, University of California. Irvine, CA; Department of Cognitive Science, University of California, San Diego, CA; Helen Wills Neuroscience Institute, University of California, Berkeley, CA; Department of Psychology, University of California, Berkeley, CA

## Abstract

Influential views of systems memory consolidation posit that the hippocampus rapidly forms representations of specific events, while neocortical networks extract regularities across events, forming the basis of schemas and semantic knowledge. Neocortical extraction of schematic memory representations is thought to occur on a protracted timescale of months, especially for information that is unrelated to prior knowledge. However, this theorized evolution of memory representations across extended timescales, and differences in the temporal dynamics of consolidation across brain regions, lack reliable empirical support. To examine the temporal dynamics of memory representations, we repeatedly exposed human participants to structured information via sequences of fractals, while undergoing longitudinal fMRI for three months. Sequence-specific activation patterns emerged in the hippocampus during the first 1-2 weeks of learning, followed one week later by high-level visual cortex, and subsequently the medial prefrontal and parietal cortices. Schematic, sequence-general representations emerged in the prefrontal cortex after 3 weeks of learning, followed by the medial temporal lobe and anterior temporal cortex. Moreover, hippocampal and most neocortical representations showed sustained rather than time-limited dynamics, suggesting that representations tend to persist across learning. These results show that specific hippocampal representations emerge early, followed by both specific and schematic representations at a gradient of timescales across hippocampal-cortical networks as learning unfolds. Thus, memory representations do not exist only in specific brain regions at a given point in time, but are simultaneously present at multiple levels of abstraction across hippocampal-cortical networks.

## Introduction

While memory for novel experiences is initially dependent on the hippocampus, neocortical regions become increasingly involved in memory retrieval over time through a process referred to as systems-level consolidation (Dudai, 2004; Frankland & Bontempi, 2005; Kumaran et al., 2016; McClelland et al., 1995; Squire et al., 1984, 2015). This change in the neural structures supporting memory occurs in a temporally protracted fashion, across months to years (Bayley et al., 2006; McClelland et al., 1995; Squire et al., 2015; Squire & Alvarez, 1995). Multiple models posit that as memories are integrated into neocortical networks, this change is accompanied by qualitative shifts in the nature of memory representations (McClelland et al., 1995; Sekeres, Winocur, & Moscovitch, 2018; Winocur et al., 2010). Specifically, consolidation is thought to aid in the extraction of common features across experiences (McClelland et al., 1995; Winocur et al., 2010), forming the basis of schemas and semantic memory in neocortical sites such as the medial prefrontal cortex (mPFC) and anterior temporal cortex, which are interconnected with the hippocampus (Gilboa & Marlatte, 2017; Preston & Eichenbaum, 2013; Ralph et al., 2017; Ranganath & Ritchey, 2012; Sekeres, Winocur, & Moscovitch, 2018; van Kesteren et al., 2012). As the transformation of specific memories into schematic representations is thought to occur slowly, frequent longitudinal sampling is required to capture the trajectory of neural representations that evolve at these protracted timescales. Consequently, there is little consensus as to how memory transformations evolve across systems consolidation, i.e. over months to years, and how the temporal dynamics of memory evolution differ across brain structures.

Evidence from human and animal lesion studies suggests that memory representations become reorganized across hippocampal-cortical networks over an extended period of at least months to years (Bayley et al., 2006; Frankland & Bontempi, 2005; Rempel-Clower et al., 1996; Scoville & Milner, 1957; Squire et al., 2015; Squire & Bayley, 2007; Zola-Morgan & Squire, 1990). According to multiple consolidation accounts, the slow extraction of information across experiences, via neocortical learning, is mediated by repeated hippocampal reactivation and hippocampal-cortical interactions. These mechanisms are thought to act as additional learning trials, allowing the neocortex to slowly discover regularities across events or individual episodes (Antony et al., 2017; McClelland et al., 1995; Sekeres, Winocur, & Moscovitch, 2018). This is especially true for cortical learning de novo (i.e. information that is not linked with prior knowledge). While cortical learning can occur at an accelerated rate when experiences are related to prior knowledge, learning of de novo representations occurs in a particularly gradual fashion (Coutanche & Thompson-Schill, 2014; Gilboa & Marlatte, 2017; Hebscher et al., 2019; Kumaran et al., 2016; McClelland, 2013; Tse et al., 2007; van Kesteren et al., 2012).

To understand the evolution of memories over time, one approach has examined how neural activity or brain-behavior relationships change with memory age (recent vs. remote memories), e.g. (Bontempi et al., 1999; J. J. Kim & Fanselow, 1992; Tallman et al., 2022). This work has suggested that the neural structures supporting memory retrieval tend to shift from the hippocampus to neocortex over time (Brodt et al., 2016; Gais et al., 2007; Gilmore et al., 2021; Smith & Squire, 2009; Takashima et al., 2006, 2009; Yamashita et al., 2009), potentially depending on the extent to which details are recovered during retrieval (Furman et al., 2012; Harand et al., 2012; Sekeres, Winocur, Moscovitch, et al., 2018). However, this approach leaves open the question of how neural representations qualitatively change, for example, from detailed and event-specific to event-general or schematic over time.

To address how representations are qualitatively transformed with consolidation, more recent work has examined the emergence of memory representations that reflect statistical regularities or structure in the environment over time after learning (Richards et al., 2014); however, the timescales of these changes, and how they differ between brain structures, remains unclear. While one study found that common neural representations across experiences emerged after 24 hours (Ritchey et al., 2015), others have found similar changes after 3 days or one week (Audrain & McAndrews, 2022; Leshinskaya et al., 2022; Tompary & Davachi, 2017), depending on when representations were measured. Specifically, one study (Tompary & Davachi, 2017) found that, after one week, representations of related items became more similar in both the hippocampus and mPFC. These findings suggest a parallel temporal development of integrated representations across the hippocampus and mPFC, contrary to consolidation accounts which juxtapose rapid hippocampal learning with slower learning in the neocortex (Frankland & Bontempi, 2005; McClelland et al., 1995). However, recent work found a distinct pattern of rapid learning of integrated representations in the medial temporal lobe followed by temporal cortex representations one week later (Leshinskaya et al., 2022). Another recent study that longitudinally tracked representations found that regularities were rapidly extracted in the hippocampus, which persisted for one week, followed by mPFC representations 2-3 weeks later (Graves et al., 2022). This pattern is consistent with proposals of slower neocortical learning, and suggests that longitudinal sampling across multiple weeks may more clearly elucidate how the dynamics of memory representations differ across hippocampal-cortical networks. Moreover, statistical learning, or the extraction of common structure across experiences, can also occur at a faster (within-session) timescale (Batterink et al., 2019; Sherman et al., 2020). These distinctive viewpoints of rapid statistical learning versus slower neocortical learning via systems consolidation further highlights the need to better understand the temporal dynamics of memory representations as learning unfolds.

Here, we examined how de novo memory representations dynamically evolve across months of learning, approaching the timescale of systems consolidation. To do so, we performed longitudinal fMRI scanning over 3 months while a small sample of participants was exposed to initially novel fractal stimuli (as part of a multi-part training study, (Miller et al., 2022)). Fractals were viewed during a serial reaction time (SRT) task, in which a subset of stimuli were embedded in temporal sequences (Fig. 1A). This allowed us to test the tenet of memory transformation accounts (McClelland et al., 1995; Sekeres, Winocur, & Moscovitch, 2018), by asking whether structured regularities across sequences developed during learning in the hippocampus and neocortex. As consolidation frameworks emphasize that multiple representations ranging in their level of detail and abstraction exist simultaneously (Gilboa & Moscovitch, 2021; Sekeres, Winocur, & Moscovitch, 2018), we tested for structured representations at multiple levels of granularity. We examined sequence-specific representations, or common patterns across fractals in the same temporal sequence, across learning (similar to pair coding neurons (Sakai & Miyashita, 1991; Schapiro et al., 2012)). We also asked whether schematic, or sequence-general representations developed by measuring representations of temporal structure that were shared between sequences.

**Figure 1.**
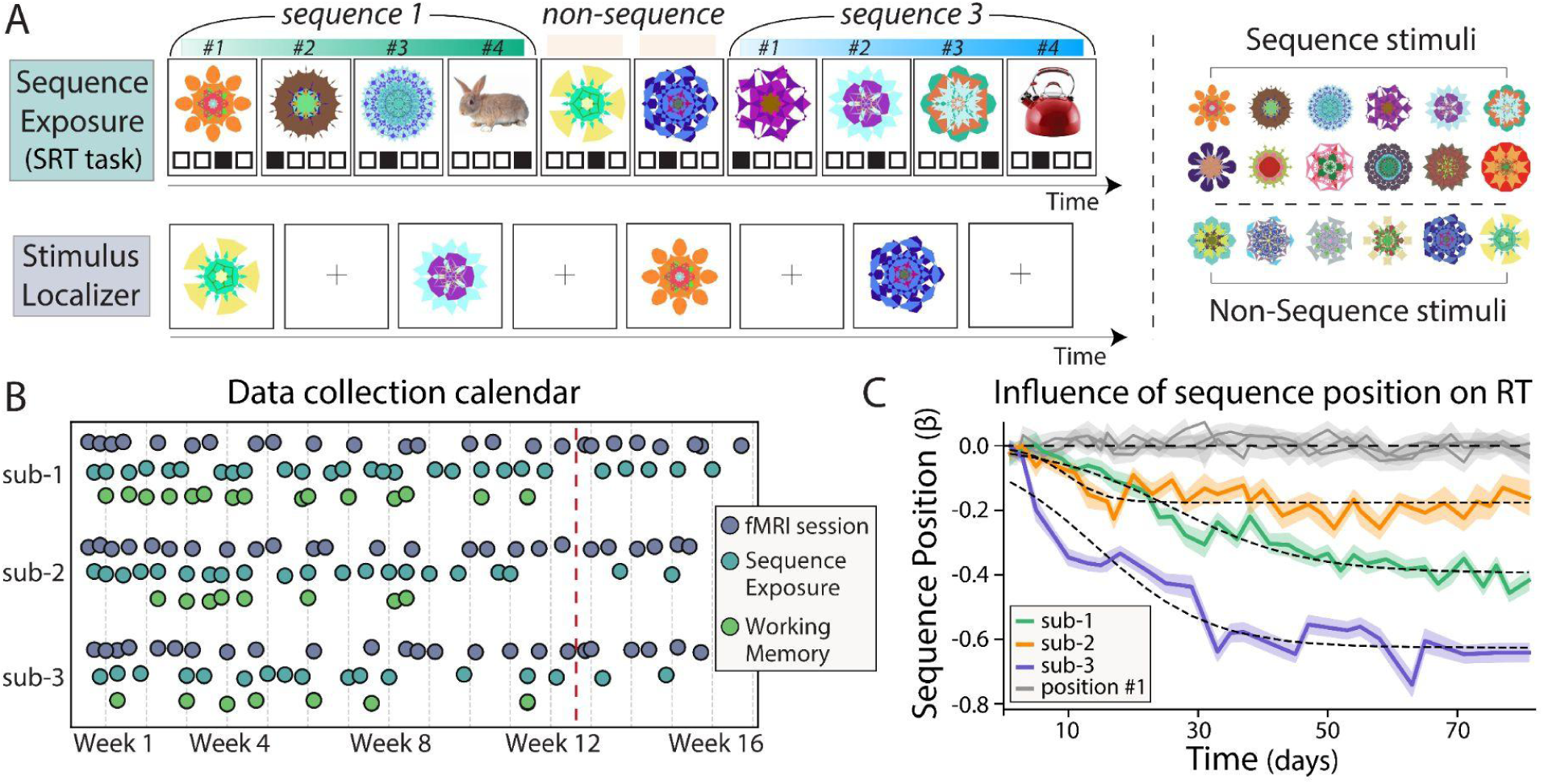
Experimental design, data collection schedule, and behavioral sequence learning. (A) Schematic of the Serial Reaction Time (SRT) task (top left), during which sequence exposure occurred. Intact sequences (three fractals followed by an object) were intermixed with non-sequence (two shown) and out-of-sequence images (not shown). Each image was associated with one of four response options, indicated by the black boxes below each image, with the correct response indicated by the filled box. The correct response was shown only during the first blocks of the first session, to facilitate stimulus-response learning. No visual indications on the screen marked sequence vs. non-sequence stimuli. Schematic of the stimulus localizer scans (bottom left), during which individual images were viewed in isolation (1.5s stimulus exposure followed by 6.5s blank screen), and participants pressed a button if a stimulus repeated. Stimulus localizer scans were used to characterize neural representations for each stimulus. An example stimulus set of one participant is shown on the right, consisting of a set of 18 fractals, 12 of which formed four temporal sequences and six non-sequence fractals. (B) Calendar of MRI (purple) and at-home sessions (SRT task, teal) for each of the three participants. During each MRI session, participants completed both the stimulus localizer scans and the SRT task (sequence exposure). The at-home training sessions consisted of modified versions of each task (Methods). The present study analyzes the first 17-18 sessions (before the red dashed line), after which new stimuli were added to the stimulus set. (C) The influence of sequence position (1-4) on response time (RT) in the SRT was assessed in each session using a general linear model, resulting in a beta coefficient for each session. Colored lines show beta coefficients averaged across sequence positions 2, 3, and 4 for each participant. Gray lines show beta coefficients for sequence position 1. Reductions in beta coefficients across time indicate that all participants showed faster RT for predicted sequence stimuli with learning. Dashed lines show logistic function fit to each participant’s data. Shaded bars indicate Standard Error of beta coefficients.

To understand the extended temporal dynamics of memory representations, we densely sampled neural representations across learning, with fMRI scans collected at least once a week for three months (Fig. 1B). This allowed us to test whether representations emerge serially or at different timescales across the hippocampus and neocortex, as predicted by rapid hippocampal versus slower cortical learning systems (Frankland & Bontempi, 2005; McClelland et al., 1995), or whether they emerge in parallel suggesting they develop from a common mechanism (e.g. as suggested by (Tompary & Davachi, 2017)). We also asked whether hippocampal representations are time-limited, consistent with a hippocampal to cortical shift predicted by ‘standard consolidation’ accounts (Alvarez & Squire, 1994; Frankland & Bontempi, 2005; Squire et al., 2015), or whether hippocampal representations persist across months of learning (Moscovitch et al., 2005; Sekeres, Winocur, & Moscovitch, 2018; Winocur et al., 2010). Finally, although consolidation accounts typically treat all neocortical representations similarly, we explored whether structured representations develop at similar or distinct timescales across neocortical structures (Sommer, 2017). To preview, our data reveal distinct neural trajectories associated with learning specific versus generalized representations, with sequence-specific representations emerging earliest in the hippocampus, followed by a cascade of cortical structures, while sequence-general representations developed first in the prefrontal cortex and then gradually in other neocortical sites and the hippocampus. Moveover, memory representations in the hippocampus and most neocortical structures tended to be sustained across learning rather than showing time-limited dynamics. Thus our findings suggest that representations tend not to be ‘transferred’ but remain present in multiple brain regions and at multiple levels of abstraction across learning and shed light on how neural representations evolve with learning.

## Results

### Behavioral evidence of sequence learning

To characterize the development of de novo memory representations, each participant was assigned a unique stimulus set of abstract fractal images, and a subset of the fractals (12/18) formed high probability sequences during a SRT task (Fig. 1A). Fractals were chosen because they have no pre-existing meaning, thus allowing us to characterize de novo learning (Naya et al., 2001; Sakai et al., 1994; Schapiro et al., 2012). Participants’ first exposure to their stimulus set occurred during the first fMRI scanning session. Here we analyze data from the 1^st^ through the 18^th^ (or 17^th^ for one participant) fMRI session, spanning approximately three months (11.5 weeks or 81 days), before additional stimuli were introduced to the stimulus set (see Methods; Fig. 1B). Participants were also exposed to the same stimuli during a single-item working memory task which was orthogonal to sequence exposure (Fig. 1B) (Miller et al., 2022). Sequence exposure in the SRT task occurred during both fMRI and at-home behavioral sessions (Fig. 1B). The SRT task allows for repeated sequence exposure and provides a clear measure of sequence learning via speeded responses to predicted sequence stimuli (those following the first item in each sequence).

We first verified that participants showed sequence learning behaviorally before examining how neural representations were shaped by sequence structure. As expected, participants showed increasingly faster response times (RTs) to predicted sequence items, providing robust evidence of sequence learning (Fig. 1C). Specifically, we assessed the influence of sequence position on RT in each behavioral and fMRI session (during intact sequence presentation, see Methods) (Bornstein & Daw, 2012). The influence of sequence position on RT for predicted images (sequence positions 2, 3, and 4) became stronger over time, as reflected in the larger negative beta coefficients that indicate faster responses (linear effect of time on RT beta coefficients: sub-1, *t*_35_ =-18.2, *p* < 10^-18^, sub-2, *t*_34_ =-5.6, *p* < 10^-5^, sub-3, *t*_25_ =-8.6, *p* < 10^-8^). In addition to examining linear changes in RT over time, we fit logistic functions which allow flexibility in when learning-related changes occur (rather than assume a constant rate of change). Significant model fits were found for each participant (cross-validated predictions, sub-1, *r* = 0.97, *p* < 10^-20^, sub-2, *r* = 0.75, *p* < 10^-7^, sub-3, *r* = 0.94, *p* < 10^-12^). As a control, we also confirmed that there was no reliable change over time in RT for the first fractal in each sequence, as it would not be predicted by preceding items (gray lines in Fig. 1C; linear effect of time: *t*_35_ = 0.49, *p* = 0.63, sub-2, *t*_34_ =-0.85, *p* = 0.40, sub-3, *t*_25_ =-0.58, *p* = 0.56; logistic function fit, sub-1, *r* = 0.23, *p* = 0.19, sub-2, *r* =-0.11, *p* = 0.65, sub-3, *r* =-0.15, *p* = 0.47). We also confirmed that responses to intact sequence fractals were faster compared to out-of-sequence presentation of the same images (Fig. S1, sub-1, *t*_36_ = 11.5, *p* < 10^-12^, sub-2, *t*_35_ = 14.7, *p* < 10^-15^, sub-3, *t*_26_ = 11.1, *p* < 10^-10^), and faster relative to non-sequence fractals (Fig. S1, sub-1, *t*_36_ = 11.2, *p* < 10^-12^, sub-2, *t*_35_ = 15.1, *p* < 10^-16^, sub-3, *t*_26_ = 10.9, *p* < 10^-10^). Together, these results indicate that responses to predicted sequence items, but not unpredicted items, became faster over time. These findings demonstrate robust evidence of sequence learning, providing a basis for examining neural representations of fractals that reflect sequence structure.

### Representational changes in the hippocampus across learning

We examined whether neural memory representations qualitatively change with learning to reflect structured regularities associated with sequence exposure. While sequence exposure occurred during the SRT task, we measured neural representations of fractals in ‘stimulus localizer’ scans, during which participants viewed each item from their stimulus set in isolation (Fig. 1A). We measured representations in the localizer scans because images were shown in rapid succession in the SRT task, making it difficult to isolate item-specific patterns that would not be systematically influenced by neighboring items. In particular, this would artificially inflate the neural similarity between items in the same temporal sequence, as these items were consistently shown together in the SRT. This approach has been extensively used in prior work to characterize how learning influences representations of items that are related in other contexts (e.g. (Deuker et al., 2016; Schapiro et al., 2012; Schlichting et al., 2015; Wammes et al., 2022)). To characterize neural representations, we extracted activation patterns during the presentation of each fractal, either within searchlight cubes or across an entire region of interest (ROI) (see Methods; Fig. 2A, left). We then measured the correlation between fractal-evoked activity patterns using a representational similarity analysis (RSA) approach (Kriegeskorte et al., 2008), resulting in a pattern similarity or correlation matrix for each fMRI session (Fig. 2A, right).

**Figure 2.**
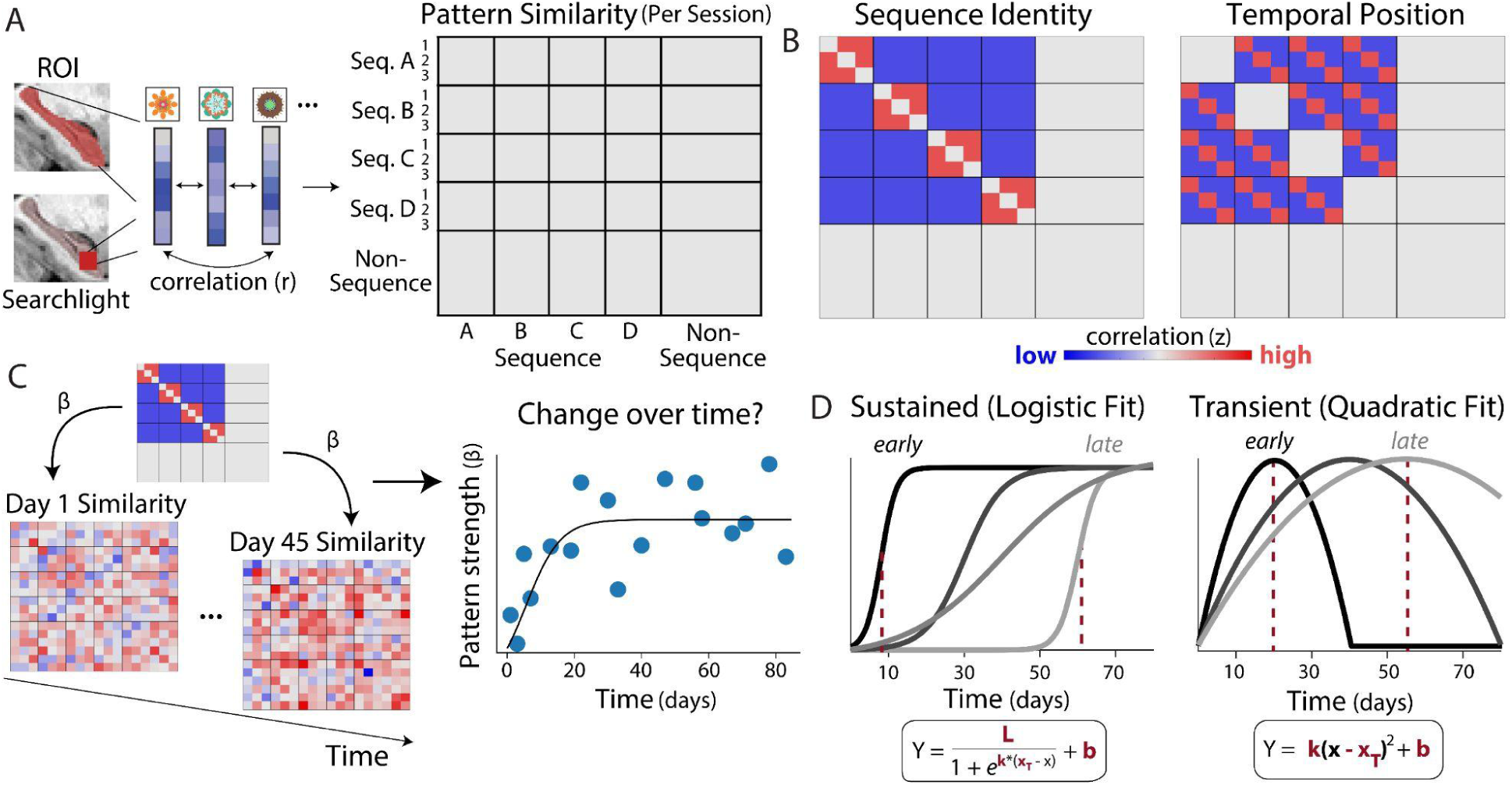
Approach for measuring representational changes across learning. (A) Pattern similarity was assessed in a given ROI or searchlight by measuring the correlation (similarity) between patterns evoked by individual fractals in the stimulus localizer scans. This resulted in a separate correlation or pattern similarity matrix for each fMRI session (right). Correlation matrices were organized by sequence identity (corresponding to four distinct sequences, A, B, C, D), ordinal position in each sequence (first, second, and third fractal), and non-sequence fractals. (B) Sequence Identity model (left) probed shared patterns across fractals in the same sequence, i.e. whether within-sequence pattern similarity (red squares) was greater than between-sequence pattern similarity (blue squares). Temporal Position model (right) probed shared patterns across fractals in the same temporal position, i.e. whether across sequence pattern similarity was greater within-position versus between-position, reflecting a sequence-general representation of temporal structure. (C) Evidence for hypothesized representations was examined using a given representation as a regressor or predictor of pattern similarity (the correlation structure) in each fMRI session, resulting in a beta coefficient reflecting ‘pattern strength’. This resulted in pattern strength values for each time point for a given searchlight or ROI (right plot) (D) Changes in pattern strength over time were assessed using two families of functions: logistic functions to model sustained or monotonically increasing representational change (left) and transient or time-limited dynamics approximated by inverted u-shaped quadratic functions (right). The free parameters for each family of functions are indicated in red.

We queried evidence for two distinct types of memory representation (Fig. 2B): 1) patterns that are shared across fractals in the same sequence, i.e. representations that reflect sequence identity (greater within vs. between sequence similarity, Fig. 2B, left) and 2) patterns that are shared between fractals in the same temporal position across sequences (greater within-position vs. between-position similarity; Fig. 2B, right), i.e. temporal representations that generalize across sequences. Sequence identity representations constitute a simple associative representation, reflecting direct item-item pairings in the SRT task. On the other hand, temporal order representations constitute schematic representations that capture generalized structure across sequences. To assess representational change during learning, we measured the presence of a given representation (e.g. sequence identity) in each fMRI session. Specifically, the strength of a representation was estimated by regressing observed pattern similarity (matrix in Fig. 2A) against a hypothesized representational structure (matrix in Fig. 2B), resulting in an estimate of ‘pattern strength’ (beta coefficient) for each fMRI session (Fig. 2C). Notably, sequence exposure in the SRT began after localizer scans in the second fMRI session, allowing pattern strength on days 1 and 2 to serve as a baseline during which no sequence-related representations should be present. We thus examined changes in pattern strength relative to these sessions when assessing learning-related representational change (Brunec et al., 2020).

After estimating pattern strength across learning, we used two primary models of temporal dynamics to capture changes in memory representations. Changes in pattern strength that were sustained across learning were probed using monotonic logistic functions (Fig. 2D, left). As depicted in Fig. 2D, logistic functions can capture changes that saturate as well as trends that are close to linear and do not reach a plateau. On the other hand, time-limited representational change was queried using inverted u-shaped quadratic functions (Fig. 2D, right). Critically, both families of functions contained free parameters that permit flexibility in temporal dynamics, such that learning-related changes can occur throughout the ∼80 day time period that we examined. To avoid overfitting, goodness of fit was estimated using cross-validation (the correlation between predicted and observed representational strength across sessions (Miller et al., 2022)).

Considering hippocampal representational change, sequence identity representations emerged over time in multivoxel patterns across the left hippocampus (Fig. 3A). Changes in sequence identity pattern strength at the ROI-level were captured by logistic functions (*r* = 0.31, *p* = 0.011), suggesting that sequence representations emerged and remained over time. The logistic function fit is shown in Fig. 3A, indicating that pattern strength changed relatively early in the experiment (inflection point = 6.5 days). In contrast, no evidence for time-limited dynamics was found in sequence identity pattern strength in the left hippocampus (quadratic model fit, 2^nd^ order coefficient =-1.8 × 10^-6^, *r* = 0.16, *p* = 0.13). While an inverted u-shaped quadratic model did not significantly predict changes in pattern strength, a linear function reliably fit the data (linear model fit, *r* =.23, *p* =.046), highlighting that a second order term did not better capture changes in pattern strength over time. In contrast to the left hippocampus, the right hippocampus did not show significant evidence of sequence identity representations (logistic fit, *r* = 0.12, *p* = 0.19, quadratic fit, 2^nd^ order coefficient = 2.2 × 10^-6^, *r* = 0.19, *p* = 0.084, indicating u-shaped change). These results suggest that left hippocampal patterns reflected sequence associations relatively early in the experiment and were sustained over time.

**Figure 3.**
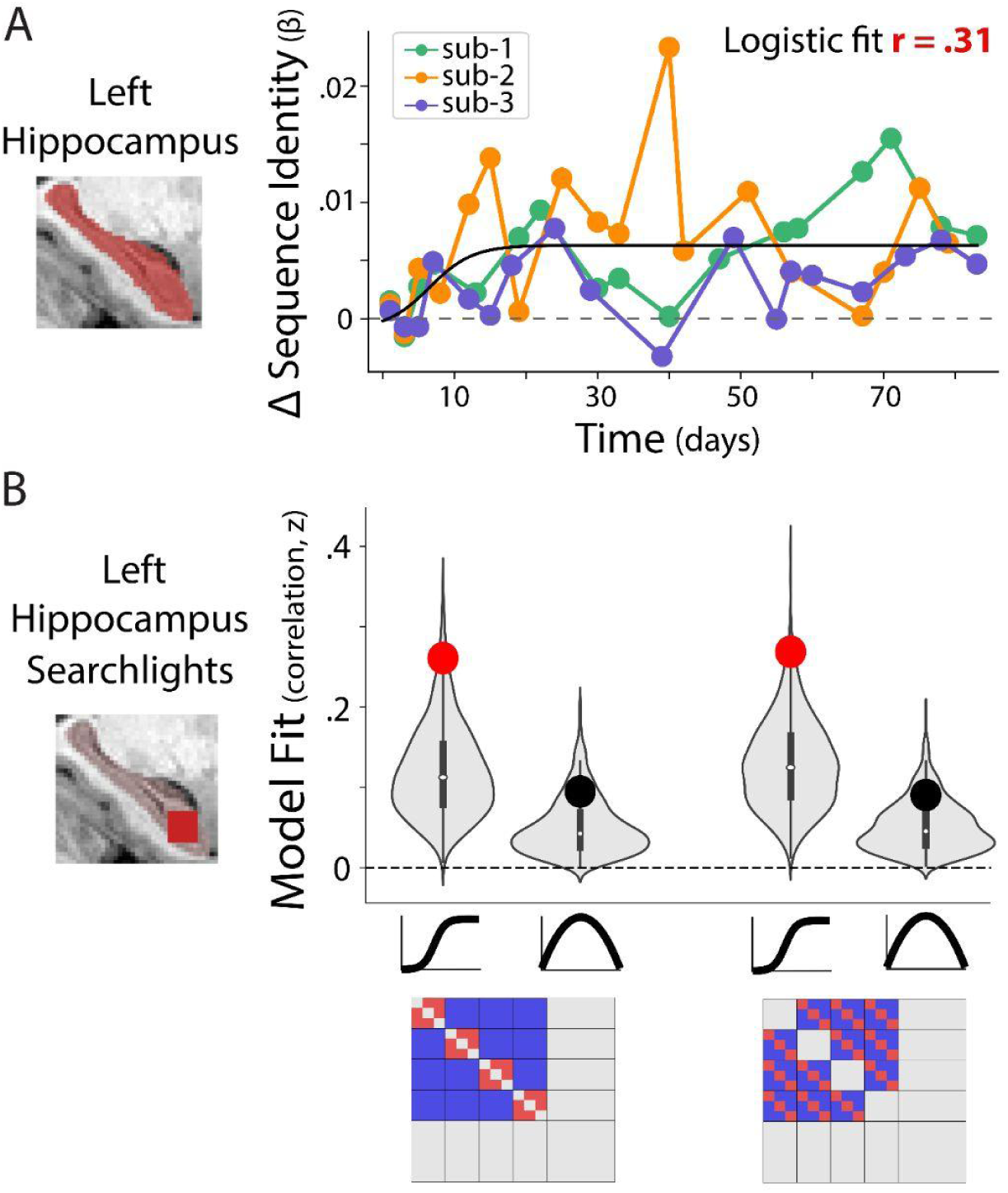
Representational change in the left hippocampus. (A) The strength of sequence identity representations across all left hippocampal voxels increased across months of learning. Data points show pattern strength (beta coefficients) for each participant relative to baseline (mean across first two sessions, prior to sequence exposure). Black line shows logistic function fit, with correlation value indicating the correspondence between predicted and actual changes in pattern strength (assessed using cross-validation). There was no reliable fit with inverted u-shaped quadratic functions. (B) Left hippocampal searchlights showed reliable development of sequence identity (left) and temporal position (right) representations indicated by logistic function fits (s-shaped curve). Model fit shows correlation between predicted and observed changes in pattern strength. Null distributions in gray show model fits obtained from permutation testing (time labels shuffled). Significant differences relative to null distributions (*p*_FDR_ <.05) indicated by red dots, non-significant differences are shown as block dots.

Turning to schematic temporal position representations, ROI analyses revealed no reliable evidence of pattern strength change across all voxels in the left hippocampus (logistic fit, *r* = 0.07, *p* = 0.32; quadratic fit, 2^nd^ order coefficient = 3.8 × 10^-6^, *r* =-0.03, *p* = 0.80) or right hippocampus (logistic fit, *r* =-0.20, *p* = 0.96; quadratic fit, 2^nd^ order coefficient =-1.0 × 10^-6^, *r* =-0.03, *p* = 0.79). This null finding was unexpected, considering extensive evidence for temporal context representations in the hippocampus (Deuker et al., 2016; Ezzyat & Davachi, 2014; MacDonald et al., 2011; Naya & Suzuki, 2011; Zou et al., 2022). However, given established functional diversity in the hippocampus (Poppenk et al., 2013; Thorp et al., 2022; S.-F. Wang et al., 2016), it is possible that schematic temporal representations may develop in more localized regions of the hippocampus (Collin et al., 2015; Dandolo & Schwabe, 2018; Sekeres, Winocur, & Moscovitch, 2018). We thus tested this possibility using a searchlight approach (J. L. S. Bellmund et al., 2022; Deuker et al., 2016; Schapiro et al., 2016). Indeed, analysis of searchlights in the left hippocampus revealed evidence for changes in the strength of temporal position representations via logistic function fits (Fig. 4B, mean logistic fit, *z* = 0.27, *p_FDR_* =.027), suggesting that more localized temporal position representations developed and persisted over time. Searchlights tended to be located in the anterior and middle portions of the hippocampus, but not the tail (Fig. S2). No reliable evidence was found for time-limited representational change in left hippocampal searchlights via inverted u-shaped quadratic fits (Fig. 3B, *z* = 0.09, *p_FDR_* =.26). Additionally, right hippocampal searchlights did not show evidence of developing temporal position representations (logistic fit, *z* = 0.12, *p_FDR_* = 0.59; quadratic fit, *z* = 0.01, *p_FDR_* =.86). These results demonstrate that sequence-general temporal position representations emerged in local patterns in the left hippocampus, which were sustained across months of learning.

**Figure 4.**
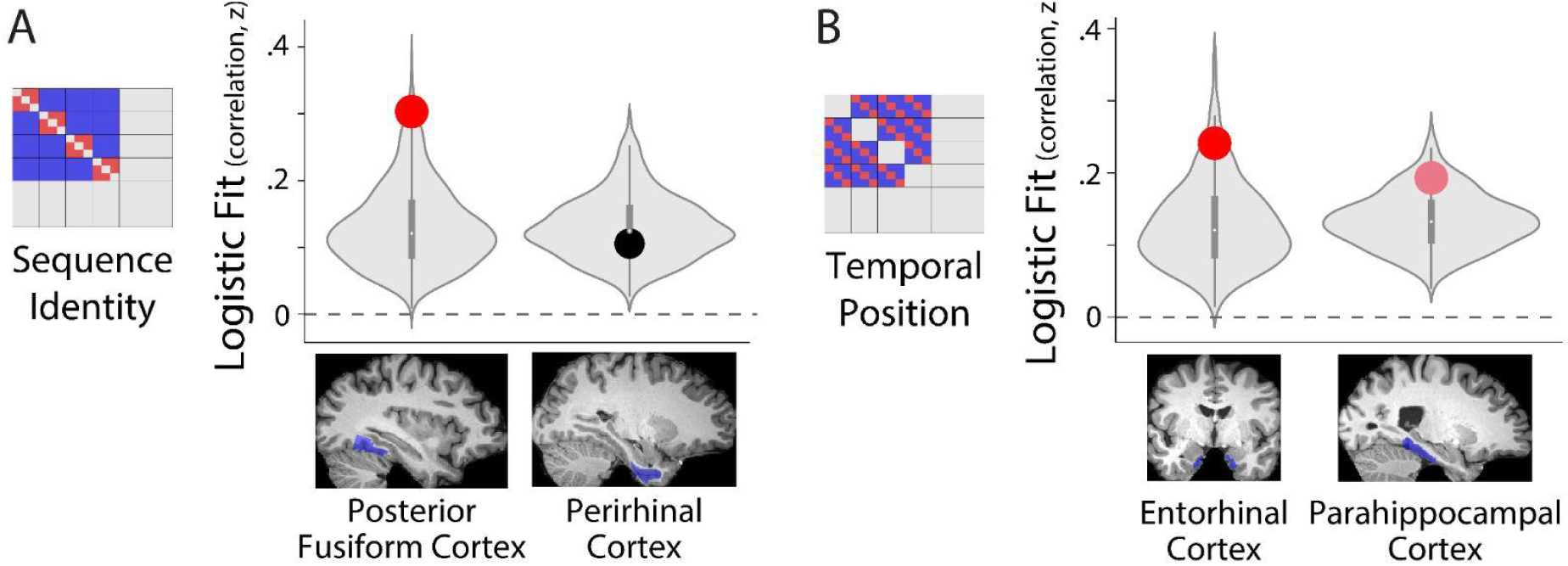
Representational changes in a priori neocortical regions across learning. (A) Sequence identity representations were examined in searchlights in the posterior fusiform cortex (left) and perirhinal cortex (right). Logistic function fits were significant in the posterior fusiform cortex, but not in perirhinal cortex. (B) Temporal position representations were probed in searchlights in entorhinal and parahippocampal cortices. Temporal position representations were significant in the entorhinal cortex and trend-level evidence was found in parahippocampal cortex. Red dots indicate significant differences from null distributions (*p*_FDR_ < 0.05); pink dots indicate trends (*p*_FDR_ < 0.10).

We next asked whether, similar to schematic temporal position representations, sequence-specific representations also developed in localized patterns within the left hippocampus (in addition to ROI-level patterns, shown above). Changes in sequence identity strength were significantly fit by logistic functions across searchlights in the left hippocampus (mean correlation, *z* = 0.25, *p_FDR_* = 0.042; Fig. 4B, left), whereas no significant evidence was found for quadratic model fits (*z* = 0.095, *p_FDR_* = 0.26). Searchlights showing sustained changes in sequence identity strength were present throughout the long-axis of the hippocampus, consistent with ROI-level results (see Fig. S2). In right hippocampal searchlights, no reliable evidence for changes in sequence identity strength over time was found (logistic fits, *z* = 0.17, *p_FDR_* = 0.19, quadratic fits, *z* = 0.05, *p_FDR_* = 0.54). These results further confirm that patterns within the left hippocampus developed sequence-specific representations that were sustained across learning.

In summary, sequence-specific and sequence-general temporal representations developed in left hippocampal patterns across months of learning. Representational patterns showed evidence for sustained but not time-limited representational change in the left hippocampus, suggesting that representations persist across months of learning, rather than being ‘transferred’ from the hippocampus as stimuli become well-learned.

### Representational changes in a priori neocortical regions across learning

Having established representational changes in the hippocampus, we examined which neocortical regions develop memory representations during learning. We first probed *a priori* neocortical regions which have established sensitivity to the content of the representations that we examined and can be readily defined based on standard anatomical or functional criterion. In particular, we expected sequence-specific patterns to emerge in regions that represent visual objects and item-item associations. We thus examined sequence-specific representations in perirhinal cortex and posterior fusiform cortex, which are responsive to visual objects and show selectivity that is shared across consistently paired items (Haskins et al., 2008; Naya et al., 2001; Sakai et al., 1994; Sakai & Miyashita, 1991; Schapiro et al., 2012). On the other hand, we hypothesized that sequence-general temporal representations would develop in regions that code for time or temporal context. Given the established sensitivity of responses in the entorhinal cortex (EC) and parahippocampal cortex (PHC) to time and temporal context (J. L. Bellmund et al., 2019; Davachi, 2006; Montchal et al., 2019; Ranganath & Ritchey, 2012; Thavabalasingam et al., 2019; Tsao et al., 2018), we tested whether temporal representations emerged in these regions. Beyond these select *a priori* regions, below we use a broader data-driven approach to isolate other neocortical sites which are likely to develop memory representations (Fig. 5). Based on the sensitivity of searchlight analyses to revealing representational change in hippocampal patterns, we primarily used searchlights for subsequent analyses (ROI-level patterns are reported to confirm null results).

**Figure 5.**
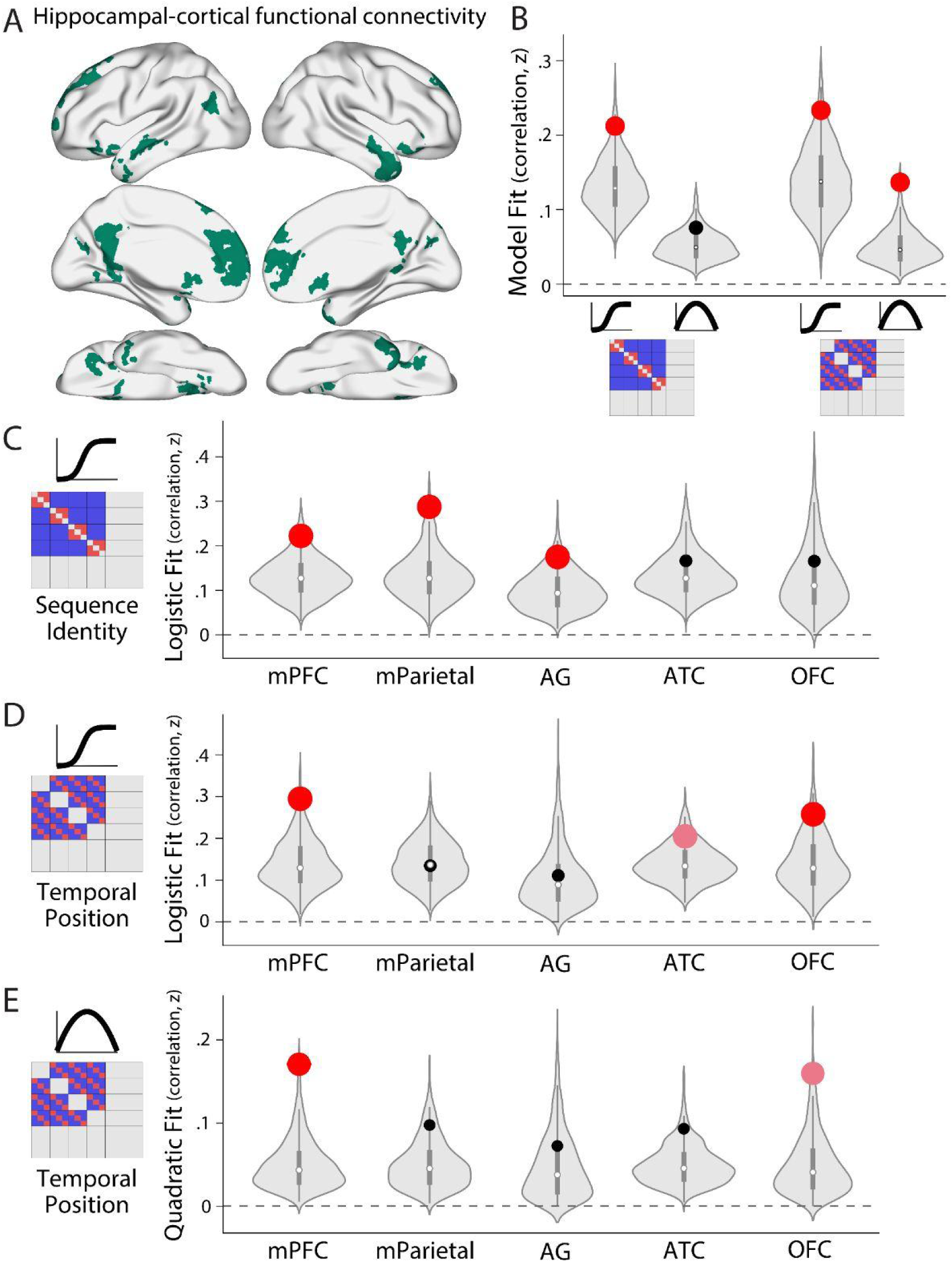
Representational changes in hippocampal-cortical networks. (A) Cortical regions showing strong connectivity with the hippocampus during sequence exposure (the SRT task) were isolated (excluding the medial temporal lobe). (B) Changes in sequence identity (left) and temporal position (right) representations were queried in searchlights across this network using logistic functions (indicated by s-shaped curve) and inverted quadratic functions (indicated by inverted u-curve). Significant logistic fits were found for both sequence identity and schematic temporal representations, indicating reliable and sustained increases in pattern strength. To localize where these representational changes occurred, the mask was subdivided into five regions. (C) Sequence identity representations reliably developed in the mPFC, medial parietal cortex (mParietal), and the angular gyrus (AG), indicated by significant logistic fits. (D) Logistic functions significantly fit the strength of temporal position representations in mPFC and OFC, with trend-level evidence in anterior temporal cortex (ATC). (E) Time-limited (inverted u-shaped) temporal position representations were found in mPFC with trend-level evidence in OFC.

We examined sequence identity representations across searchlights in the perirhinal and posterior fusiform cortices. Logistic function fits indicated significant increases over time in the strength of sequence identity representations in posterior fusiform cortex searchlights (Fig. 4A, *z* = 0.30, *p_FDR_* = 0.027) but not in the perirhinal cortex searchlights (Fig. 4A, logistic fits, *p_FDR_* = 0.34). The lack of reliable representational change in perirhinal cortex was confirmed by examining patterns across the perirhinal cortex ROI instead of in searchlights (right perirhinal cortex, logistic fit, *r* = 0.16, *p* = 0.13; left perirhinal cortex, logistic fit, *r* = 0.17, *p* = 0.12). No evidence for reliable quadratic changes over time was found in either region (posterior fusiform searchlights, *z* = 0.06, *p_FDR_*= 0.48, perirhinal searchlights, *z* = 0.04, *p_FDR_*= 0.67, right perirhinal quadratic fit, 2^nd^ order coefficient =-2.1 × 10^-6^, *r* =-0.08, *p* = 0.73, left perirhinal quadratic fit, 2^nd^ order coefficient = 4.9 × 10^-6^, *r* = 0.02, *p* = 0.44). These results indicate that, in addition to the left hippocampus, patterns reflecting sequence associations emerged in the posterior fusiform cortex and were sustained across learning. However, there was no evidence for shared sequence representations in the perirhinal cortex.

The entorhinal cortex (EC) and parahippocampal cortex (PHC) encode elements of time and context (J. L. Bellmund et al., 2019; Davachi, 2006; Montchal et al., 2019; Ranganath & Ritchey, 2012; Thavabalasingam et al., 2019; Tsao et al., 2018). We thus investigated whether generalized representations of temporal position developed in the EC and PHC across learning. Indeed, temporal position representations emerged over time in searchlights in the entorhinal cortex, as indicated by significant logistic fits (Fig. 4B, *z* = 0.24, *p_FDR_*= 0.044) with trend-level evidence for logistic fits in parahippocampal cortex searchlights (Fig. 4B, *z* = 0.19, *p_FDR_* = 0.062). In contrast, no evidence of time-limited representational change was found in these regions (EC, *z* = 0.046, *p_FDR_* = 0.58; PHC, *z* = 0.054, *p_FDR_* = 0.57). These findings indicate that generalized or schematic representations of temporal structure develop and remain sustained in entorhinal and parahippocampal cortices.

### Representational changes in hippocampal-cortical networks across learning

Beyond the above a priori neocortical regions, we sought to uncover representational change in canonical hippocampal-cortical networks (Ranganath & Ritchey, 2012; Ritchey & Cooper, 2020; Vincent et al., 2006), including sites such as the mPFC and anterior temporal cortex, which are likely to contain schematic and semantic memory representations (Gilboa & Marlatte, 2017; Gilboa & Moscovitch, 2021; Sekeres, Winocur, & Moscovitch, 2018). As there is no standard approach to *a priori* define where memory representations should emerge within these regions, we used a data-driven approach to isolate neocortical sites showing functional connectivity with the hippocampus during the SRT task (Fig. 5A, medial temporal lobe cortex was excluded as it was queried in prior analyses, see Methods). When examining searchlights across this network of cortical regions (Fig. 5A), both sequence identity and schematic temporal position representations emerged in a temporally sustained manner, as evidenced by significant logistic function fits (Fig. 5B, sequence identity, *z* = 0.21, *p_FDR_* = 0.042, temporal position, *z* = 0.23, *p_FDR_* = 0.042). Evidence for time-limited temporal position representations was also found, as indicated by reliable quadratic fits (Fig. 5B, *z* = 0.14, *p_FDR_* = 0.028); however quadratic fits of sequence identity strength were not significant (Fig. 5B, *z* = 0.08, *p_FDR_* = 0.26).

To localize representational changes, we divided the neocortical structures shown in Fig. 5A into five ROIs (mPFC, medial parietal cortex, angular gyrus, anterior temporal cortex, and orbitofrontal cortex). Changes in sequence identity pattern strength (Fig. 5C), as evidenced by logistic function fits, were found in searchlights in the mPFC (*z* = 0.22, *p_FDR_*= 0.042), medial parietal cortex (*z* = 0.29, *p_FDR_*= 0.027), and angular gyrus (*z* = 0.18, *p_FDR_*= 0.044), with no reliable changes in anterior temporal (*z* = 0.17, *p_FDR_* = 0.23) or orbitofrontal cortex searchlights (OFC, *z* = 0.17, *p_FDR_* = 0.23). No reliable evidence for time-limited sequence identity representations were found in searchlights in the five ROIs examined (quadratic function fits, *p_FDR_*> 0.16). These results indicate that sequence-specific representations emerged and were sustained over time in the mPFC, medial parietal cortex, and angular gyrus.

Turning to schematic temporal position representations (Fig. 5D-E), significant representational change evidenced by logistic function fits (Fig. 5D) were found in searchlights in the mPFC (*z* = 0.29, *p_FDR_* = 0.027) and OFC (*z* = 0.26, *p_FDR_* = 0.044), with a trend in anterior temporal cortex (*z* = 0.20, *p_FDR_* = 0.062). Logistic function fits were not reliable in medial parietal cortex (*z* = 0.13, *p_FDR_* = 0.52) or angular gyrus searchlights (*z* = 0.11, *p_FDR_* = 0.36). Inverted u-shaped functions (Fig. 5E) reliably fit changes in temporal position representations in mPFC (*z* = 0.17, *p_FDR_*= 0.028) and OFC searchlights (*z* = 0.16, *p_FDR_*= 0.062), but not in other regions (*p_FDR_* > 0.16). These results indicate that schematic position representations developed in mPFC, OFC, and anterior temporal cortex, with evidence for both temporally sustained and time-limited representations in the prefrontal cortex (mPFC, OFC).

To better understand the nature of changes in mPFC and OFC temporal position representations, which were fit by both logistic and inverted u-shaped functions, we asked whether searchlights fit by these functions were spatially distinct or overlapping. Distinct populations of voxels would suggest a diversity of temporal dynamics within a region, while highly overlapping searchlights suggest that temporal dynamics may resemble a mixture of logistic and quadratic dynamics. In the mPFC, logistic versus quadratic searchlights were largely non-overlapping in two of three participants, but were almost completely overlapping in one participant. In the OFC, one participant showed highly overlapping logistic and quadratic searchlights, with one participant showing only significant logistic searchlights, and the other participant showing only significant quadratic searchlights. Thus, there is evidence for both distinct populations of voxels that change with different temporal profiles within mPFC and OFC, but also some evidence for a mixture of dynamics. Below, we further examine the temporal dynamics of memory representations in these regions.

In summary, these findings indicate that multiple regions across hippocampal-cortical networks developed structured sequence-specific and sequence-general representations across learning. Multiple temporal dynamics were observed, such that sequence-specific representations showed evidence of sustained changes over time, while schematic representations showed both time-limited and temporally sustained dynamics.

### Timescales of representational change

After isolating hippocampal and neocortical sites that developed structured memory representations across learning, we compared the timescale of changes across brain regions and distinct memory representations. A major prediction of consolidation accounts (McClelland et al., 1995) is that hippocampal representations should emerge prior to neocortical representations (schematized in Fig. 6A). To test this idea, we estimated *when* the greatest extent of representational change occurred by examining the inflection points of fitted logistic and quadratic functions (in searchlights with significant fits, see Methods). We also visualized logistic and quadratic function fits (when they reliably fit the data) to better understand the temporal dynamics of representational change across months of learning.

**Figure 6.**
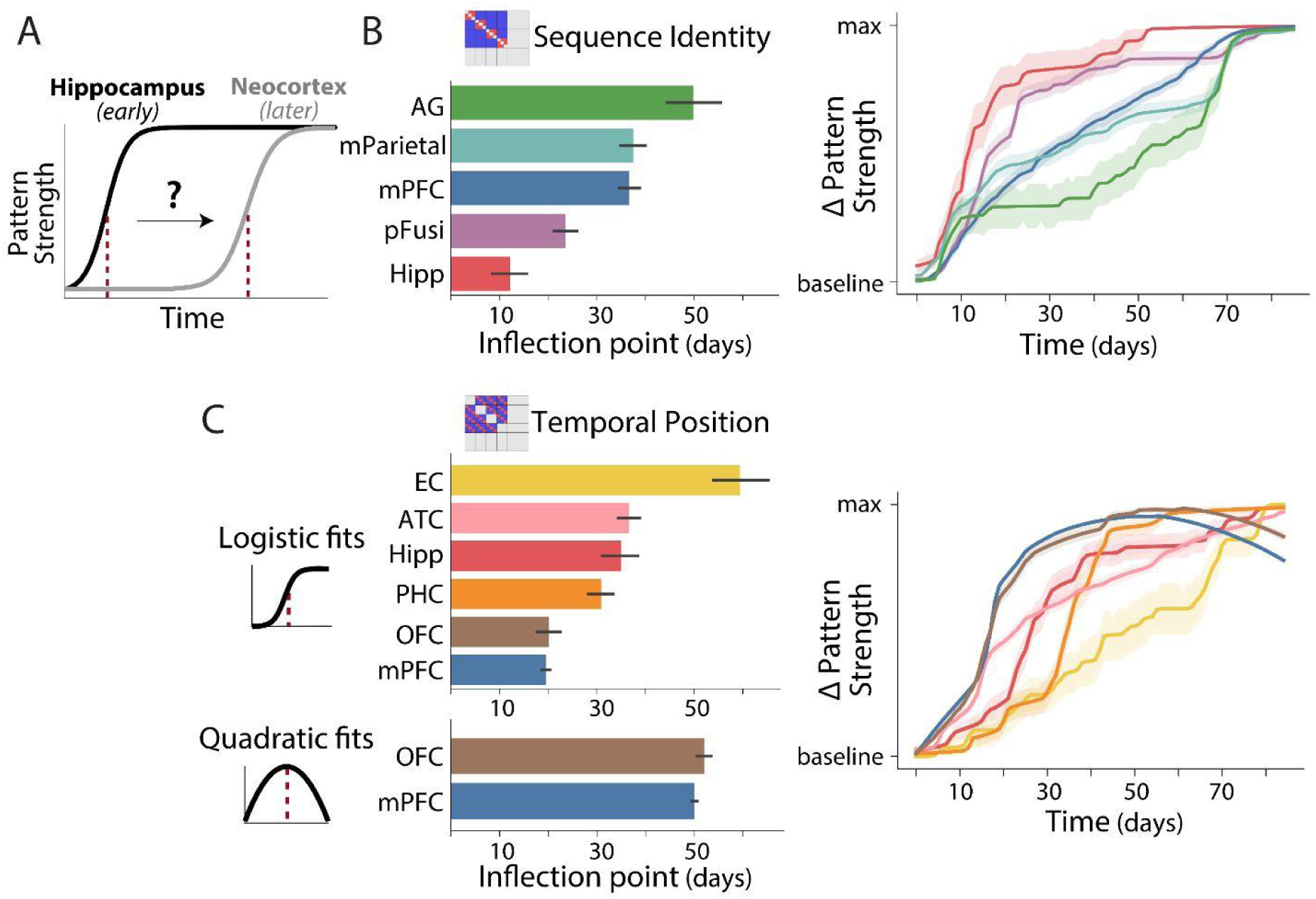
Timescales of representational changes. (A) Hypothesized early emergence of representational changes in the hippocampus versus neocortex, which should manifest as earlier inflection points of logistic functions fits (dotted red lines). (B) Timescales of changes in sequence identity representations. Left plot shows the average inflection point of logistic function fits in regions that showed significant changes in sequence identity representations (Figs. 4, 5). Inflection points were pooled across significant searchlights in each brain region. Regions are ordered by the mean inflection point from bottom to top. Right plot visualizes changes over time, or average idealized pattern strength over time across significant searchlights. Pattern strength values were normalized to fixed levels to facilitate comparison of timescales across regions. (C) Timescales of changes in schematic temporal position representations, following the same conventions as (B). Timescales of changes are shown separately for regions showing logistic changes (top) and for regions showing inverted u-shaped temporal dynamics, pooled across searchlights that were significantly fit by inverted quadratic functions (bottom). Inflection points for quadratic fits indicate when representational strength was at peak levels and began to decline. Right plot shows idealized pattern strength changes over time, relative to baseline levels across logistic and quadratic changes. Error bars on all plots indicate bootstrapped 68% confidence intervals across searchlights.

We first examined the temporal dynamics of sequence-specific representations within searchlights across the hippocampus and neocortical regions. Searchlights in the left hippocampus showed the earliest development of sequence identity representations (indicated by the smallest inflection point, mean = 12.2 days), followed by searchlights in the posterior fusiform cortex (mean = 23.5 days), mPFC (mean = 36.6 days), medial parietal cortex (mean = 37.5 days), and angular gyrus (mean = 49.9 days, Fig. 6B, left). Reliable differences across brain regions were found using a nonparametric analysis of variance (*p* < 10^-5^; Methods). The inflection point in hippocampal searchlights was reliably earlier than all cortical regions (vs. posterior fusiform, *p* = 0.005; vs. mPFC, *p* < 10^-5^; vs. medial parietal cortex, *p* = 2×10^-5^; vs. angular gyrus, *p* < 10^-5^). Sequence identity representations in posterior fusiform cortex emerged earlier than those in mPFC (*p* = 0.00012) and parietal cortex (vs. medial parietal cortex, *p* = 0.00016; vs. angular gyrus, *p* = 3×10^-5^). Sequence representations developed latest in the angular gyrus (vs. mPFC, *p* = 0.017, vs. medial parietal cortex, *p* = 0.04). To visualize the timescale of changes across regions, we plotted the idealized logistic function fits for each region (Fig. 6B, right, averaged across searchlights). Here the progression of representational change can be observed, emerging earliest in the hippocampus (red), followed by high-level visual cortex (purple), to medial prefrontal and parietal cortices (blues), and the angular gyrus (green).

Turning to the dynamics of schematic temporal position representations, as we found that both logistic and inverted u-shaped functions significantly fit changes over time, we examined the inflection points of each family of functions separately (Fig. 6C, left). The inflection points of logistic functions (Fig. 6C, top left) significantly varied across brain regions (*p* < 10^-5^). The earliest inflection points were found in mPFC (mean = 19.6 days) and OFC searchlights (mean = 20.2 days; Fig. 6C). Inflection points in parahippocampal cortex (PHC), left hippocampus, anterior temporal cortex, and entorhinal cortex (EC) searchlights lagged behind those in mPFC (vs. PHC, *p* = 0.0004; vs. hippocampus, anterior temporal cortex, EC, all *p* < 10^-5^) and OFC (vs. PHC, *p* =.0088; vs. hippocampus, *p* =.00025; vs. anterior temporal cortex, *p* =.00093; vs. EC, *p* < 10^-5^). Surprisingly, the EC showed the latest development of temporal position representations (Fig. 6C; mean = 59.4 days, greater than all other regions, *p*’s <.0008). These results indicate that schematic representations first developed in the mPFC and OFC, followed by a gradual emergence of representations in the MTL and anterior temporal cortex.

When examining time-limited or inverted u-shaped changes in schematic representations (Fig. 6C, bottom left), inflection points occurred relatively late in learning (mean = 50 days for mPFC searchlights, 52 days for OFC searchlights). Inverted u-shaped model fits are shown in Fig. S3, indicating an increase in strength until ∼7 weeks, when representations began to decline. Pattern strength did not return to baseline levels after three months. This temporal profile is compatible with logistic function fits, which increased during a similar timeframe of 6-7 weeks before reaching a plateau (Fig. S3). Therefore, when visualizing representational change (Fig. 6C, right), we summarized the best-fit dynamics across logistic and inverted u-shaped functions for PFC regions (see Methods). These results suggest early increases in schematic temporal representations in the PFC, followed by a partial decline later in learning.

Across all regions showing sequence and temporal position representations (Fig. 6B vs. 6C), temporal position representations tended to emerge later than sequence identity (mean difference in inflection points of logistic functions = 5.2 days, *p* =.002). The earliest inflection points for schematic temporal position representations in mPFC and OFC were significantly later than left hippocampal inflection points for sequence identity representations (mean difference vs. mPFC = 7.4 days, *p* = 0.00036; difference vs. OFC = 8 days, *p* = 0.02). This supports the notion that sequence associations were first learned in the hippocampus, followed by extraction of sequence-general representations in the PFC.

### Trade-offs between the expression of specific vs. schematic sequence representations

The prior results indicate the presence of at least two types of memory representations, sequence-specific and sequence-general representations, across multiple brain regions and in some cases the same region (see summary of representational changes in Fig. 7). The existence of both specific and generalized representations, in response to the same item cues, raises the question of whether specific and schematic representations were active in mind simultaneously, or whether there are trade-offs between the relative dominance of each representation (illustrated in Fig. 8A). In other words, the expression of sequence-specific information may trade-off with sequence-general representations, and vice versa. To test this idea, we assessed the trial-by-trial strength of each representation (see Methods). We adopted an informational connectivity approach (Anzellotti & Coutanche, 2018) to examine: 1) whether the expression of sequence-specific representations was negatively correlated with sequence-general representations across trials (suggesting a tradeoff between representations) and 2) whether each type of representation was positively correlated between brain regions (suggesting coherent expression of a given representation across regions at any time point). To examine learning-related changes, we compared informational connectivity at the beginning and end of learning.

**Figure 7.**
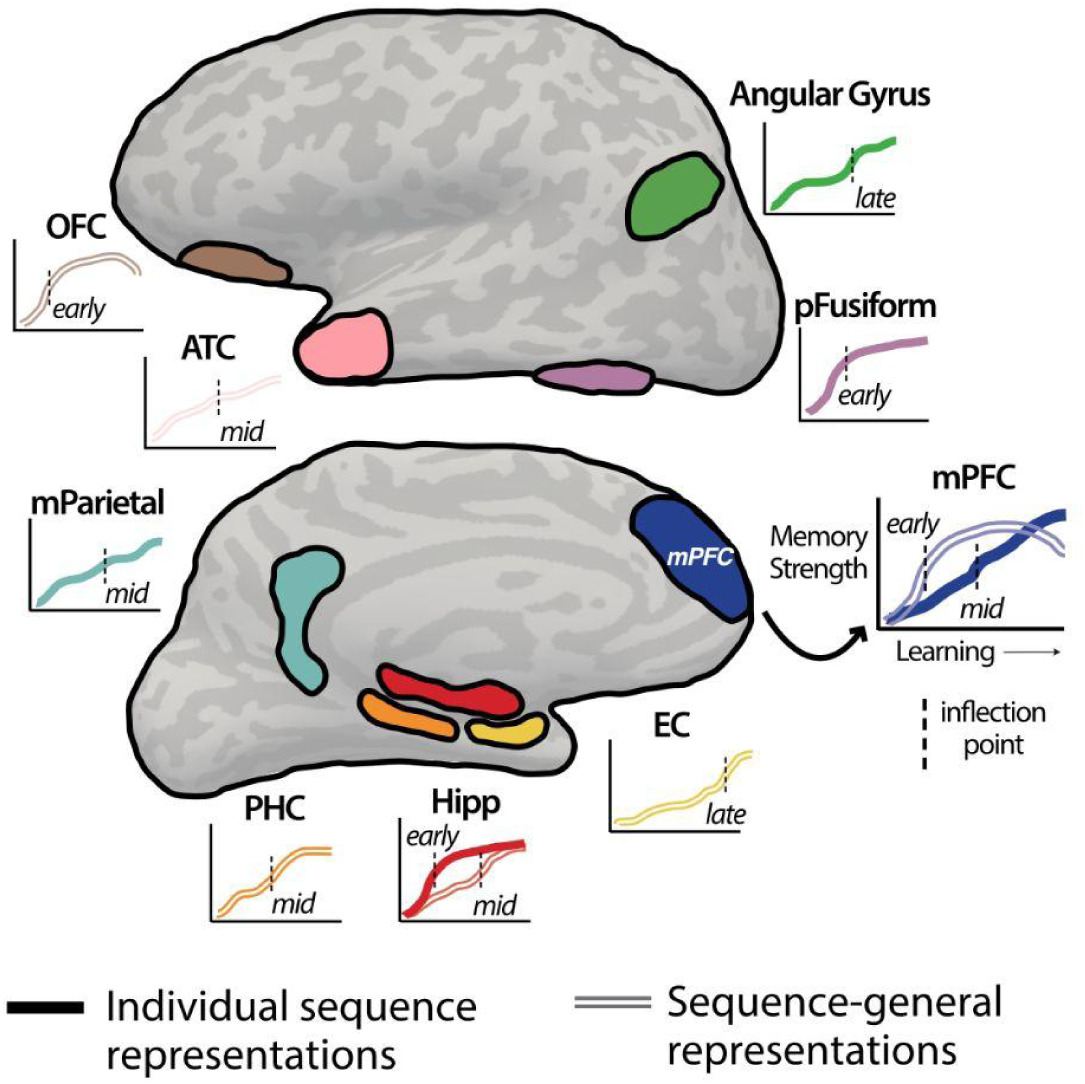
Schematic of temporal dynamics of memory representations across regions. Brain regions showing significant development of memory representations over time are highlighted, with approximated changes in representational strength shown for each region. As indicated for the mPFC, x-axis depicts time across learning and y-axis indicates changes in representational strength. Black dotted line on each plot depicts the approximate inflection point of fit logistic functions. Sequence identity representations are shown in full (bolded) line, and schematic, sequence-general representations are shown in striped line. Labels on each plot indicate approximately when representations developed, early, middle (‘mid’), or late during the three month learning period.

**Figure 8.**
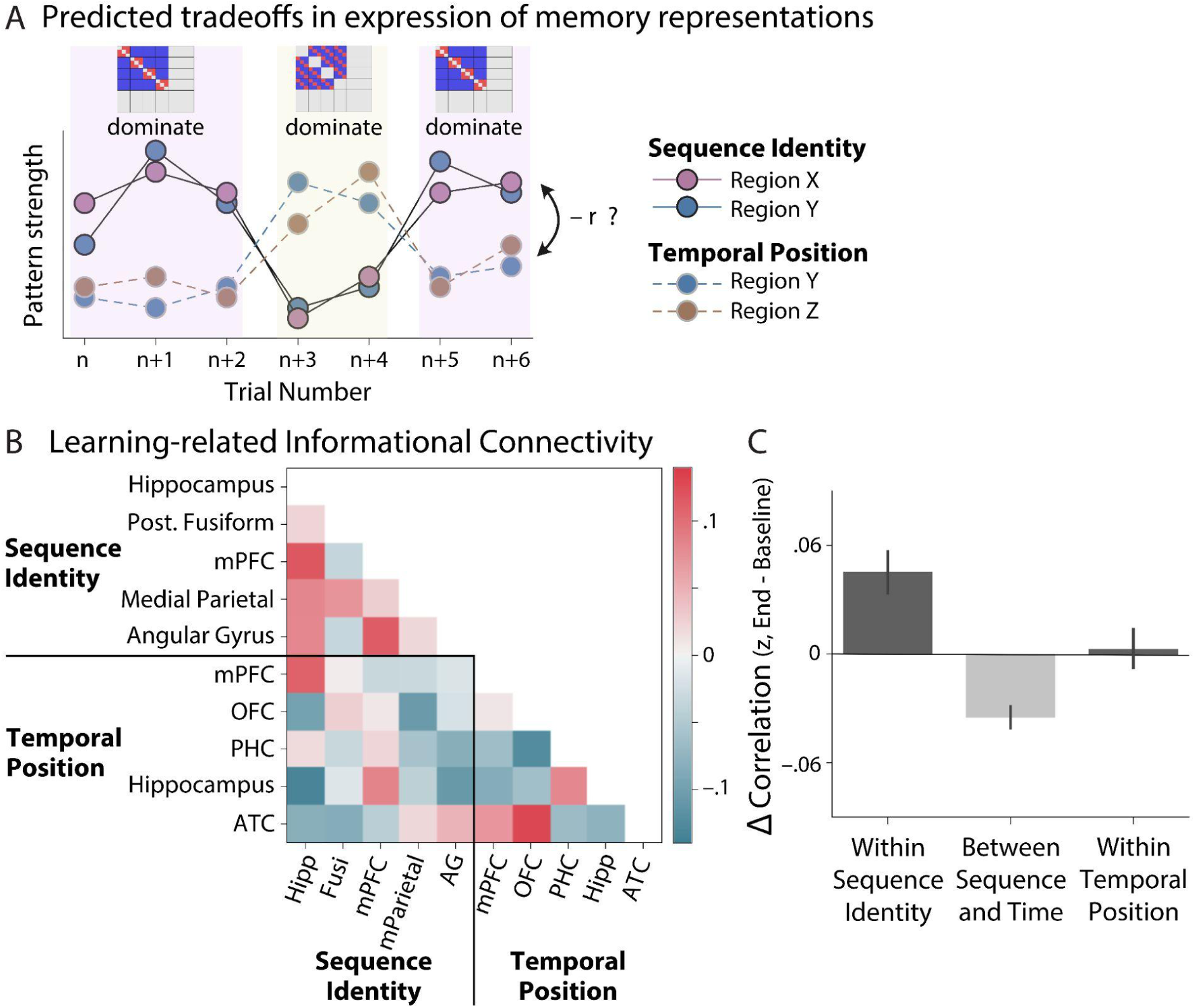
Tradeoffs in the expression of distinct memory representations at short timescales. (A) Hypothesized fluctuations in the trial-by-trial expression of memory representations. We expected tradeoffs or negative correlations between fluctuations in pattern strength between sequence-specific vs. sequence-general representations across trials, suggesting time periods in which one representation is dominantly expressed. For a given representation, e.g. sequence identity, we expected pattern strength would be correlated across brain regions. (B) Learning-related changes in informational connectivity across all brain regions showing representational change (excluding entorhinal cortex). The correlation in pattern strength was examined after learning (sessions 10 and beyond, after ∼36 days), relative to before sequence exposure (days 1 and 2), yielding learning-related changes in informational connectivity (positive values in red, negative values in blue). The correlation matrix is ordered by representation and brain region, black lines mark divisions between within-and between-representation correlations. (C) Average changes in learning-related informational connectivity, for sequence information (left bar), between sequence and temporal position information (middle bar), and for temporal position information (right bar).

Learning-related changes in informational connectivity across memory representations and brain regions is shown in Fig. 8B. While some brain regions (and representations) showed increases in informational connectivity (indicated by red), others showed decreases (blue values), suggesting distinct expression of memory representations across trials. As predicted, when examining informational connectivity across sequence identity and temporal position representations, correlations in the trial-by-trial strength of these representations decreased across learning (lower left portion of matrix in Fig. 7B, middle bar in Fig. 8C; *t*_13_ =-5.07, *p* = 0.0002). On the other hand, coherent expression of sequence identity representations, indicated by increases in trial-by-trial correlations of pattern strength, was found for sequence identity representations (top left portion of matrix in Fig. 8B, left bar in Fig. 8C; *t*_13_ = 3.70, *p* = 0.0024). However, unlike sequence identity, temporal position representations did not appear to show systematic changes in coherence across all region pairs (lower right portion of matrix in Fig. 8B, right bar in Fig. 8C; *t*_13_ = 0.23, *p* = 0.82). These findings suggest that specific vs. generalized memory representations were expressed in a temporally segregated manner, and that sequence-specific representations were expressed in a coherent fashion across hippocampal-cortical networks.

## Discussion

It is widely accepted that memories are reorganized across hippocampal-cortical networks, over an extended timescale of months to years. This process is thought to support the extraction of common features across experiences, forming the basis of schemas and semantic memory. Despite these established views, empirical tests of the extended temporal dynamics over which memory representations develop, and how they differ between brain regions, have been rare. To understand the temporal evolution of memory representations, we performed longitudinal fMRI across three months of an extended learning experience. During learning, initially novel fractal stimuli were shown in temporal sequences, allowing us to examine how sequence structures influenced memory representations. Activation patterns in the hippocampus and multiple neocortical regions evolved to reflect both sequence-specific and schematic, sequence-general relationships between fractals. Sequence-specific patterns developed most rapidly in the left hippocampus, emerging in the first 1-2 weeks of learning. After the hippocampus, neocortical sequence-specific representations emerged in a gradual progression from high-level visual cortex, to medial prefrontal and parietal cortex, and lastly in the angular gyrus across multiple weeks. Schematic, sequence-general representations of temporal position, on the other hand, showed distinct temporal dynamics, developing earliest in the PFC (∼3 weeks), followed by the parahippocampal cortex, hippocampus, anterior temporal cortex, and latest in the entorhinal cortex. Most sequence-specific and schematic representations tended to remain sustained across months of learning, as opposed to showing transient, time-limited dynamics. We also found that the expression of specific versus schematic memory representations tended to trade-off across trials, suggesting that representations at a particular level of abstraction were dominant at a given moment. Together, these findings indicate that memory representations emerge at a gradient of timescales across hippocampal-cortical networks, with hippocampal patterns emerging earliest. Rather than ‘transferring’ between brain regions or representational formats, memory representations were simultaneously present in multiple regions and levels of abstraction across learning.

Several systems consolidation frameworks contrast fast hippocampal acquisition with slower learning rates in the neocortex, such that the hippocampus slowly ‘teaches’ other structures about recently acquired memories through repeated reactivation and interactions (Frankland & Bontempi, 2005; Klinzing et al., 2019; Kumaran et al., 2016; McClelland et al., 1995; Sekeres, Winocur, & Moscovitch, 2018; Winocur & Moscovitch, 2011). Consistent with this idea, we found that the left hippocampus showed the earliest development of individual sequence representations out of the regions that we examined, with neocortical representations emerging later. This temporal separation of memory representations supports the notion of rapid hippocampal versus slower neocortical learning, and shows that this distinction is evident when extensively sampling memory representations across months of learning. However, we found that schematic representations emerged earliest in the prefrontal cortex and later in the hippocampus (and MTL cortex). While the overall pattern of results support faster hippocampal versus slower neocortical learning, this finding indicates that the hippocampus (and MTL) does not always show the most rapid formation of memory representations and highlights the role of the prefrontal cortex in schematic representations (see further discussion below).

Sequence representations emerged in the posterior fusiform cortex, a high-level visual object processing region, approximately one week after the hippocampus. Post-encoding interactions between the hippocampus and unimodal sensory association areas (overlapping and adjacent portions of the lateral occipital complex and fusiform gyrus) have been shown to support later memory (de Voogd et al., 2016; Kark & Kensinger, 2019; Liu et al., 2018; Murty et al., 2017; Schlichting & Preston, 2014; Tambini et al., 2010; Tambini & D’Esposito, 2020). Combined with these prior findings, our results suggest that representations are first reorganized across the hippocampus and high-level sensory association cortices, and that interactions between these regions are likely to mediate initial systems memory reorganization. These interactions may continue during early weeks of visual learning in parallel with changes in memory representations (c.f. (Lewis et al., 2009)). Posterior fusiform cortex representations were followed by the medial prefrontal and parietal cortex, and finally by the angular gyrus latest in learning. This temporal pattern of development further suggests that representations in multimodal association cortices, at least for novel stimuli, may be learned via interactions with both the hippocampus and sensory association cortices.

Most systems consolidation frameworks do not differentiate between neocortical sites when considering memory reorganization across hippocampal-cortical networks (McClelland et al., 1995; Nadel et al., 2000; Squire et al., 2015; Winocur & Moscovitch, 2011). While some consolidation frameworks propose that different neocortical regions support distinct memory representations (Moscovitch & Gilboa, 2021; Sekeres, Winocur, & Moscovitch, 2018) (see also (Ranganath & Ritchey, 2012; Ritchey & Cooper, 2020)), the question of whether learning rates or timescales differ across neocortical sites is not typically addressed. One recent framework (Kaefer et al., 2022) proposes that memory reorganization temporally progresses along the principal connectivity gradient (Huntenburg et al., 2018; Margulies et al., 2016), such that reactivation and reorganization first propagates from the hippocampus to transmodal default mode network (DMN) regions (which are most strongly connected to the hippocampus), and later to primary sensory cortices, although this has not yet been empirically tested. Our finding that sequence-specific representations emerged in high-level visual cortex prior to DMN regions (mPFC, medial parietal cortex, and angular gyrus) instead suggests that, at least for de novo visual learning, memory reorganization may initially occur via hippocampal interactions with sensory association cortices. It is possible that distinct learning paradigms would result in different temporal dynamics; for example, cross-modal learning with direct links to prior knowledge would more strongly engage the mPFC and other DMN regions (Liu et al., 2017; van Kesteren et al., 2013; X. Wang et al., 2020) and may result in earlier memory reorganization in the DMN. Regardless, our findings indicate a gradient in the development of memory representations across neocortical sites, rather than a unified emergence of neocortical learning after the hippocampus.

In addition to sequence-specific patterns, schematic or sequence-general representations of temporal position emerged in multiple sites. Unlike sequence-specific patterns which first emerged in the hippocampus, schematic representations developed earliest in the prefrontal cortex (mPFC and OFC). This highlights the role of the prefrontal cortex in extracting schematic memory representations by combining information across associations. These results are consistent with the role of the mPFC in schema processing and representations, integration, and memory consolidation (Audrain & McAndrews, 2022; Baldassano et al., 2018; Gilboa & Marlatte, 2017; Giuliano et al., 2021; Preston & Eichenbaum, 2013; Tompary & Davachi, 2017; van Kesteren et al., 2012; Zheng et al., 2021) as well as schematic and abstract state space representations in the OFC (Boorman et al., 2021; Mızrak et al., 2021; Schuck et al., 2016; Wilson et al., 2014; Zhou et al., 2021) and recent evidence that it supports temporal memory (Johnson et al., 2022). Moreover, the delayed emergence of schematic representations in the MTL and anterior temporal cortex relative to the prefrontal cortex, combined with a partial decline of prefrontal representations late in learning, suggest that schematic representations may potentially migrate from the PFC to portions of temporal cortex over time. This has also been suggested in a prior training study, which found that mPFC activation declined between 1 and 3 months as activation in the anterior temporal cortex increased (Sommer, 2017). Additionally, the observation of delayed schematic representations in the anterior temporal cortex resonates with prior work showing categorical representations in this area months after training (Ghazizadeh et al., 2018) and its role in semantic and conceptual knowledge (Ralph et al., 2017; Yee et al., 2014) and long-term retention of semantic information (Holdstock et al., 2002). In summary, these results highlight that schematic representations first emerge within nodes of the default (mPFC) and limbic (OFC) networks (Yeo et al., 2011) and continue to develop across other sites in these networks over time (hippocampus, MTL cortex, and anterior temporal cortex).

Early formulations of systems consolidation (Squire, 1992; Squire & Alvarez, 1995) postulate that memories are transferred from the hippocampus to more stable neocortical sites over time, based on observations of temporally graded retrograde amnesia after hippocampal damage (Anagnostaras et al., 1999; Scoville & Milner, 1957; Squire & Bayley, 2007; Zola-Morgan & Squire, 1990). However, the extent to which the hippocampus continues to be involved in memory over time has been extensively debated (e.g. (Gilmore et al., 2021; Nadel & Moscovitch, 1997; Tallman et al., 2022)). Hippocampal activation during retrieval has been shown to decline as memories become more remote (Gais et al., 2007; Gilmore et al., 2021; Smith & Squire, 2009; Takashima et al., 2006, 2009), but this pattern is not always observed and may reflect changes in the quality of retrieved content over time (Ezzyat et al., 2018; Furman et al., 2012; Gilboa et al., 2004; Harand et al., 2012; Sekeres, Winocur, Moscovitch, et al., 2018; Viskontas et al., 2009). Here we measured the temporal dynamics of structured representations over multiple months of continued exposure, as a model for the formation of new schematic memories. If representations become independent of the hippocampus as they are learned across repeated exposures, this predicts that hippocampal representations should decline within three months of learning. We found no evidence for time-limited dynamics of hippocampal representations, but instead found that representations appeared sustained. This suggests that both specific and generalized hippocampal representations persist after information is well-learned, rather than being ‘transferred’ or declining as neocortical representations develop. Prior observations of hippocampal activation declining with long-term learning (Sommer, 2017) suggest that activation levels may not always track the presence of structured memory representations. It may be the case that we observed sustained hippocampal representations because participants were continuously exposed to sequences across learning. However, it seems unlikely that repeated exposure would continually drive reconsolidation processes and trigger new hippocampal encoding, given the absence of any explicit novelty or changes in stimulus contingencies during learning (Sinclair et al., 2021; Sinclair & Barense, 2018, 2019). Moreover, prior observations of distinct hippocampal representations for 10-12 year old memories (Bonnici et al., 2013; Bonnici & Maguire, 2018), generalized representations from one week (Tompary & Davachi, 2017) to a month (Dandolo & Schwabe, 2018) after learning, as well as conceptual representations (Morton et al., 2021; Quian Quiroga et al., 2009) suggest that intensive exposure is not required to maintain hippocampal representations, corroborating our finding that at least some forms of hippocampal representations persist over time.

In contrast to views that a memory representation resides in a single location or small set of regions, it is likely that memories are represented in multiple formats or levels of abstraction and more than one brain region simultaneously (Gilboa & Moscovitch, 2021; Roy et al., 2022; Sekeres, Winocur, & Moscovitch, 2018). This notion is supported by our finding that sequence-specific and generalized representations emerged across multiple brain regions during learning. This viewpoint raises the possibility that representations of varying levels of granularity, such as detailed versus abstract representations, are expressed to different degrees depending on task demands, as shown in recent behavioral studies (Tompary et al., 2020; Zeng et al., 2021). Using an informational connectivity approach (Anzellotti & Coutanche, 2018), we found that the trial-by-trial correlation between the strength of sequence-specific versus sequence-general representations decreased from before to after learning. This suggests that the expression of distinct versus generalized representations becomes segregated on a moment-to-moment basis, and that detailed versus abstract representations may dominate at different times (c.f. (Goshen et al., 2011)). However, because memory representations were characterized here in the absence of retrieval demands (i.e., during the localizer task), we were unable to link fluctuations in the expression of distinct representations with ongoing behavior.

A large body of prior work has shown that hippocampal and neocortical memory representations change at varying timescales, including within single sessions, over one day, and across multiple days, depending on when neural measures were collected (e.g. (Brodt et al., 2018; Cowan et al., 2021; Dandolo & Schwabe, 2018; Fernandez et al., 2023; Leshinskaya et al., 2022; Ritchey et al., 2015; Schapiro et al., 2012, 2013; Schlichting et al., 2015; Tompary & Davachi, 2017)). Our findings extend this work by showing that memory representations continue to develop across multiple months of learning, and that the timescales over which representations evolve differs across brain regions and levels of abstraction. However, the ability to resolve rapid versus long timescale changes is limited in this dataset, as it is difficult to assess the reliability of change between individual data points with a small sample size. Instead, we analyzed systematic changes across 17-18 time points per participant, to track the evolution of representations across an extended window of months. This design was chosen to minimize data loss and ensure that participants would remain committed throughout the study. Limited sampling of few individuals likely results in findings that do not necessarily generalize to the population of healthy adults. Therefore, given the small sample, instead of examining variance between participants, when possible, we combined data across participants using fixed-effects analyses, allowing us to make the strongest inference in the current sample (as opposed to the general population, see (Fries & Maris, 2021; Vezoli et al., 2021)). Similar designs that extensively sample a small group of individuals have also been used across multiple fMRI studies, such that sufficient statistical power is obtained by extensively sampling individuals rather than sampling less frequently in a smaller population (Gordon et al., 2017; Gratton & Braga, 2021; Kragel et al., 2021; Laumann et al., 2015; Miller et al., 2022; Naselaris et al., 2021; Nee, 2019; Newbold et al., 2020; Popham et al., 2021). Moreover, the collection of multiple data points over time is critical to characterize memory processes that unfold at extended timescales (Miller et al., 2022; Vanasse et al., 2022; Zou et al., 2022).

In conclusion, by tracking fMRI activation during months of learning, we found that de novo memory representations emerged at a gradient of timescales across hippocampal-neocortical networks. Although our findings support the notion of faster hippocampal versus slower neocortical learning, they also demonstrate that there is not a simple mapping between learning rates and brain regions, as distinct temporal dynamics were found for specific versus generalized representations. We found little evidence for a transfer of representations between the hippocampus and neocortex, as predicted by some consolidation accounts. Instead, our data show that memory representations are multiplexed, existing simultaneously across distinct levels of abstraction and multiple brain structures at a given time, which are differentially expressed on a moment-to-moment basis. These findings shed light on how memory representations evolve across extended timescales in hippocampal-neocortical networks. Moreover, the current approach of tracking memory over time within individuals may be fruitful for targeting and modulating subject-specific memory representations.

## Methods

### Participants

Data were collected from three healthy, adult volunteers. Because of the large amount of data collected and intensive nature of the design, all participants were members of the research team who completed the study over the same time period. One participant was a 34-year-old female (sub-001), one was a 25-year-old male (sub-002), and one was a 37-year-old female (sub-003). The University of California, Berkeley Committee for the Protection of Human Subjects (CPHS) approved the study protocol and no participants reported any contraindications for MRI.

### Experimental Design

To adequately sample and characterize the timescale of changes in memory representations, we collected longitudinal fMRI and behavioral data over multiple sessions across four months of learning (Fig. 1B). Four fMRI sessions were collected during the first week of the experiment to ensure sampling of initial learning, followed by 1-2 fMRI sessions per week (depending on participant and MRI scanner availability). FMRI sessions consisted of rest scans, stimulus localizer scans, the SRT task, and a Working Memory (WM) task. The SRT task and stimulus localizer scans are described in detail below; rest scans and the WM task were not analyzed in the current work. Changes in prefrontal activity and representations during the WM task have been previously described (Miller et al., 2022).

To regularly expose participants to the stimuli and SRT task, at-home behavioral sessions were interspersed with fMRI sessions. During behavioral sessions, participants either performed the SRT task or a Working Memory (WM) task (Fig. 1B). The at-home SRT task was administered in order to avoid gaps of more than three days without exposure to the task. After the primary portion of the experiment, or approximately 81 days (11.5 weeks, 17 fMRI sessions for one participant and 18 for two participants, indicated by the red line in Fig. 1B), new stimuli were introduced to examine the influence of prior knowledge on subsequent learning. Here, we report data from the first 17 or 18 sessions to characterize initial learning.

### Data and code availability

All neuroimaging data are openly available in the Brain Imaging Data Structure format (Gorgolewski et al., 2016) (BIDS) on OpenNeuro: https://openneuro.org/datasets/ds003659 (Markiewicz et al., 2021). Analysis code will be made available on Open Science Framework.

### Stimuli

To study de novo learning, each participant was assigned a unique stimulus set of 18 novel fractal images. Each image was an algorithmically-generated fractal consisting of multiple colors, and the 18 images for each participant were balanced according to the primary color group of each image (determined using a k-means clustering algorithm on each fractal in the *sklearn* package: https://scikit-learn.org/). These fractals were chosen because they are visually complex, approximately uniform in size, cannot be easily verbalized, have no pre-existing meaning, and thus have been used in prior work to study learning-related changes in neural representations (H. F. Kim et al., 2015; Sakai & Miyashita, 1991; Schapiro et al., 2012).

Because the study participants were also on the research team, we avoided participants gaining any foreknowledge of their stimulus set by generating thousands of images and randomly selecting each training set from among these images. Thus, each participants’ first exposure to their training set occurred during the localizer scans in the first fMRI session. The unique 18 stimuli for each participant were then used in all of the following fMRI and behavioral training sessions. Of the 18 fractal stimuli in each participant’s training set, 12 were randomly assigned to be part of four sequences in the SRT task, with each sequence consisting of three fractals and an object image. Half of the object images were animals and the other half were tools. In order to characterize de novo learning, responses to object images were not characterized in the current study.

### Serial Reaction Time (SRT) Task

Participants performed a SRT task to repeatedly expose them to structured experience (statistical regularities) in the form of regularly occurring temporal sequences. During this task, participants made button presses in response to each stimulus. The stimulus set consisted of each participant’s set of 18 fractal stimuli and six objects (three animals and three tools) for a total of 24 stimuli. The SRT task consisted of three phases: an initial phase in which stimulus-response mappings were learned, a second phase during which stimulus sequences were present, and a third phase in which new stimuli were introduced. Here, we analyze behavioral data from the first two phases of the task (before new stimuli were introduced) to characterize initial sequence learning, and hippocampal functional connectivity in the same sessions. However, note that changes in fMRI activation associated with fractals was assessed in the localizer scans (see below) and not during the SRT task.

The first phase of SRT task was implemented in the first two sessions of the study (one fMRI session followed by an at-home behavioral session) during which participants were trained to criterion to associate each of the stimuli with one of four button press responses. Participants were first exposed to their stimulus set during their first scanning session. During every block of the SRT task, each of the 24 stimuli were shown once in a randomized order. Each fMRI run consisted of six SRT task blocks (six presentations of each stimulus). No sequence information was present during the first two sessions (first fMRI and first behavioral session). Each stimulus was presented on the screen for 2.3 seconds (followed by a blank screen of 0.7 s between stimuli) with four response options shown as black squares below the stimulus (corresponding to the middle finger of the left hand, ring finger of the left hand, ring finger of the right hand, and middle finger of the right hand). During the first two blocks of the first scanning session, the correct response was highlighted (square corresponding to the response was shown in red instead of black) to allow participants to view the correct response and facilitate learning. Then, participants completed 10 more blocks during which the correct response was not shown but feedback was provided for 200 ms (when a correct response was made the square turned blue, but incorrect responses were indicated by the selected option turning red). After the first scanning session, participants performed an at-home session to ensure the learning of stimulus-response mappings. Participants completed a minimum of five blocks of the task, and continued until a criterion of 80% accuracy at the item-level was reached (>=80% of correct first responses for all stimuli across all blocks; 7-15 blocks of training were required to reach criterion). The stimulus-response mappings remained constant throughout the study.

After the completion of training to criterion, temporal sequences of stimuli were embedded in the SRT task, beginning in the second fMRI session (after the stimulus localizer scans). Of the 24 trained stimuli (18 fractals and six objects), 16 stimuli were assigned to form four distinct sequences, with each sequence containing three fractals followed by an object (**Fig. 1A**). Although sequences were not explicitly instructed, all participants had knowledge of the sequence manipulation; thus, reductions in response time for sequence stimuli likely reflect both explicit and implicit learning. As in the initial phase of this task, each stimulus was shown once during each block (set of 24 trials) and the four response options were indicated below the stimulus as four black squares. Participants were instructed to press the appropriate button for each stimulus. Each stimulus was shown for 1.95 s (fMRI sessions) or 1.8 s (behavioral sessions) followed by a blank screen for 400 ms. Sequences were presented in a probabilistic manner, such that three of the four sequences were presented in an intact fashion in each block and each sequence was intact on 75% of blocks in each session (i.e. in 12/16 blocks during fMRI sessions). In each block, the order of the presentation of stimuli was randomized with the exception of the presentation of the three intact sequences. Stimuli from the non-intact sequence (one sequence per block) were presented in a random order with the stipulation that at least two stimuli separated the non-intact sequence stimuli. Feedback was provided throughout the experiment as described above in the training to criterion phase. The fMRI sessions contained 18 blocks of the SRT task and the at-home behavioral sessions consisted of 26 blocks. Stimuli were presented in a randomized order (no sequence information was present) during the first two blocks of each session which served to acclimate participants to the task.

### Analysis of response time (RT) in the SRT task

To verify the presence of sequence learning in the SRT task over time, we 1) calculated general linear models (GLMs) in each behavioral and fMRI session to estimate the influence of sequence position on RT and 2) compared RTs for intact sequence presentation versus out-of-sequence presentation of the same images and non-sequence images. In the first approach, RT for all trials in each session were modeled with the following regressors (Bornstein & Daw, 2012): 24 regressors coding for the presence of each unique stimulus to capture stimulus-specific RTs, four regressors each coding for sequence position (1, 2, 3, or 4) when intact sequences were presented, and nuisance regressors (one coding for cumulative trial number in the session, one coding for the presence of a repeated motor response). Trials with incorrect responses or long RTs (> 2 standard deviations from the mean of each session) were excluded from all analyses. This allowed us to estimate the influence of sequence position on RT, resulting in a parameter estimate (beta coefficient) for each sequence position and session. For simplicity, parameter estimates were averaged across positions 2-4 in each session (as shown in Fig. 1). To quantify changes across learning, linear regressions with time were performed, in addition to fitting logistic functions to data points over time using cross-validation (using the same methods described below in <u>Modeling changes in representational strength across learning</u>).

To verify faster responses to intact sequence presentation, we compared average RT across sessions for intact sequences, out-of-order (shuffled) presentation of the same sequence stimuli, as well as non-sequence images (shown in Fig. S1). As in the prior analyses, RTs for sequence positions 2-4 were averaged before comparing RTs across conditions. For each participant, paired t-tests were used to test for significance in RT differences between trial types across sessions.

### Stimulus localizer task scans

To characterize change in neural activity associated with learning, participants were exposed to stimuli from the SRT task during separate ‘localizer’ scans. Here, stimulus presentation was separated in time to isolate item-specific patterns (1.5 s stimulus presentation followed by a blank screen for 6.5 s) (Visser et al., 2016; Zeithamova et al., 2017). Sequence stimuli were not presented consecutively to avoid inflating the similarity of responses to fractals in the same temporal sequence due to temporal autocorrelation. During the task, participants were instructed to view each image and perform a 1-back task (make a button press when an image was presented two times in a row). No demands were present to overtly retrieve any information from the SRT task (e.g. button press associated with each image, or whether an item was part of a temporal sequence).

Each localizer scan consisted of two blocks; during each block all images in each participant’s stimulus set (fractals and objects) were shown once with two images repeated consecutively per block (repeated images were chosen randomly). Stimuli were shown in a randomized order in each block, with the stipulation that at least two stimuli separated the presentation of images belonging to the same temporal sequence. Each fMRI session contained four localizer scans, with a total of eight repetitions of each item in a participant’s stimulus set.

### Object-selective category functional localizer task

To localize object processing regions in ventral temporal cortex (posterior fusiform cortex), standard category-level functional localizer scans were collected during two fMRI sessions for each participant (separate sessions from the main study). These scans were collected after sessions 1 and 5 for sub-1, sessions 1 and 15 for sub-2, and sessions 5 and 14 for sub-3. Participants performed a one-back task while viewing blocks of animals, tools, objects, faces, scenes, and scrambled images. All images were presented on phase scrambled backgrounds. Each block lasted for 16 s and contained 20 stimuli per block (300 ms stimulus presentation followed by a blank 500 ms inter-stimulus interval). Two stimuli were repeated in each block and participants were instructed to respond to stimulus repetitions via button press. Each scan (three scans per session) contained four blocks of each stimulus class, which were interleaved with five blocks of passive fixation.

### fMRI acquisition

All neuroimaging data were collected on a 3 Tesla Siemens MRI scanner at the UC Berkeley Henry H. Wheeler Jr. Brain Imaging Center (BIC). Whole-brain Blood Oxygen Level-Dependent (BOLD) fMRI (T_2*_-weighted) scans were acquired with a 32-channel RF head coil using a 2x accelerated multiband echo-planar imaging (EPI) sequence [repetition time (TR) = 2 s, echo time = 30.2 ms, flip angle (FA) = 80°, 2.5 mm isotropic voxels, 52 slices, matrix size = 84 × 84]. Anatomical MRI scans were collected at two timepoints across the study and registered and averaged together before further preprocessing. Each T_1_-weighted anatomical MRI was collected with a 32-channel head coil using an MPRAGE gradient-echo sequence [repetition time (TR) = 2.3 s, echo time = 3 ms, 1 mm isotropic voxels]. For each scan, participants wore custom-fitted headcases (caseforge.com) to facilitate a consistent imaging slice prescription across sessions and to minimize head motion during data acquisition (Power et al., 2019).

In each 2-hr scanning session (except for the first session), participants completed the following BOLD fMRI scans: (1) 9 min eyes-closed rest run, (2) three 9 min runs of the stimulus localizer task, (3) three 6 min runs of the SRT task, (4) 9 min eyes-closed rest block, (5) 9-min stimulus localizer block, (6) four 6 min runs of the WM task. During the first fMRI session, participants completed two 7.6 min runs of the SRT task (first phase of SRT, in which no sequences were present). The present work focuses on the stimulus localizer scans and SRT task. See (Miller et al., 2022) for descriptions of changes in WM behavior and PFC delay period activity across learning.

### MRI preprocessing and general analysis methods

Preprocessing of the neuroimaging data was performed using fMRIPrep version 1.4.0 (Esteban et al., 2019), a Nipype (Gorgolewski et al., 2018) based tool. Each T1w (T1-weighted) volume was corrected for INU (intensity non-uniformity) using *N4BiasFieldCorrection* v2.1.0 (Tustison et al., 2010) and skull-stripped using *antsBrainExtraction.sh* v2.1.0 (using the OASIS template). Brain tissue segmentation of cerebrospinal fluid (CSF), white-matter (WM) and gray-matter (GM) was performed on the brain-extracted T1w using fast (Zhang et al., 2001) (FSL v5.0.9).

Functional data was slice time corrected using 3dTshift from AFNI v16.2.07 (Cox, 1996) and motion corrected using mcflirt (Jenkinson et al., 2002) (FSL v5.0.9). This was followed by co-registration to the corresponding T1w using boundary-based registration (Greve & Fischl, 2009) with 9 degrees of freedom, using flirt (FSL). Motion correcting transformations and BOLD-to-T1w transformation were concatenated and applied in a single step using *antsApplyTransforms* (ANTs v2.1.0) using Lanczos interpolation. Framewise displacement (FD) was calculated for each functional run, using their implementations in Nipype (following the definitions by (Power et al., 2014)). A set of physiological regressors were extracted to allow for anatomical component-based noise correction (aCompCor, (Behzadi et al., 2007)). Principal components are estimated after high-pass filtering the preprocessed BOLD time-series (using a discrete cosine filter with 128s cut-off). Anatomical CompCor components are calculated within the intersection of a whole-brain mask and the union of CSF and WM masks calculated in T1w space, after their projection to the native space of each functional run (using the inverse BOLD-to-T1w transformation). Many internal operations of FMRIPREP use Nilearn (Abraham et al., 2014), principally within the BOLD-processing workflow. For more details of the pipeline see https://fmriprep.readthedocs.io/en/latest/workflows.html. Spatial smoothing was performed using a 4mm full-width half maximum kernel in SPM12.

A group-level anatomical template was created using Advanced Normalization Tools (ANTs, antsMultivariateTemplateConstruction2) (Avants et al., 2011) using the skull-stripped T1-weighted scans of all three participants. This template was used for group-level analysis of hippocampal-cortical functional connectivity during the SRT (Fig. 5), otherwise all analyses were performed in ROIs in subject-specific space.

Several nuisance regressors were included in general linear models (GLMs) when modeling BOLD fMRI data (stimulus localizer scans and category-level functional localizers). Nuisance regressors consisted of: six motion parameter (MP) estimates, their first temporal derivative, four regressors each indicating the first four volumes of the scan prior to task onset (to account for T1 equilibration), regressors indicating high motion time points (all volumes with FD >.5 mm, including one volume before and two after), the top eight aCompCor principal components extracted from fMRIprep (white matter and cerebrospinal fluid signals), and low frequency trends (cutoff frequency =.0078 Hz or 128s).

Neuroimaging files were loaded and operated on using the Nibabel *(Brett et al., 2020)* and *Nilearn* packages (https://nilearn.github.io/; (Abraham et al., 2014)). For all plots, error bands reflect bootstrapped 68% confidence intervals as implemented in the *Seaborn* package (Waskom et al., 2020) (approximately the standard error of the mean).

### Region-of-Interest (ROI) definition

Medial temporal lobe (MTL) regions (hippocampus, entorhinal cortex, perirhinal cortex, and parahippocampal cortex) were defined anatomically based on each participant’s T1 weighted scan. The hippocampus was defined using FSL’s FIRST (Patenaude et al., 2011) followed by manual corrections to ensure accurate definitions according to (Pruessner et al., 2000). Entorhinal, perirhinal, and parahippocampal cortices were manually defined according to (Moore et al., 2014). Searchlights were further used as a data-driven approach to examine localized representations in the hippocampus, as we did not have sufficient spatial resolution to delineate hippocampal subfields and exploratory analyses suggested that sequence representations did not clearly correspond to common definitions of the anterior vs. posterior hippocampus across all participants. The searchlight approach allowed us to identify sites of representational change in each participant while permitting variability in the precise location of changes across participants.

As in prior work, posterior fusiform cortex was functionally defined from category-level functional localizer scans. Category-selective localizer scans were modeled with 16 s boxcar regressors for each stimulus category (objects, phase scrambled images, faces, and scenes) along with nuisance regressors (see <u>MRI</u> <u>processing and general analysis methods</u> above). Object processing regions were localized from the contrast of objects > scrambled images at P <.0001 and posterior fusiform cortex was defined by selecting voxels at this threshold that were located in fusiform cortex (Schwarzlose et al., 2008).

Regions showing reliable functional connectivity with the hippocampus during the SRT were isolated to probe cortical memory representations. The bilateral hippocampus was used as a seed region, and correlation maps (Fisher Z-transformed) for each session were transformed into ANTs template space (see below for details, <u>Hippocampal functional connectivity during SRT task</u>). A t-test was performed across all sessions and subjects, and a strict threshold of *t* > 17 (50 voxel extent) was used to isolate the canonical set of regions that typically interact with the hippocampus (Ranganath & Ritchey, 2012). The threshold of *t* > 17 was not set a priori, but was chosen as it resulted in a clean separation of cortical regions (more liberal thresholds resulted in activation that spread across distinct anatomical regions, e.g. anterior temporal cortex and OFC). In order to examine regions that were not part of the MTL (captured in anatomical ROI analyses), voxels in the hippocampus and surrounding MTL were manually removed. The mask was then subdivided into cortical ROIs: mPFC, medial parietal cortex, anterior temporal cortex (bilateral), OFC (bilateral) and the left angular gyrus.

### Hippocampal functional connectivity during SRT task

Correlations or functional connectivity (FC) with the hippocampus (bilateral) was measured during the SRT task to isolate cortical regions which may develop memory representations over time. To do so, low-frequency trends were removed from the preprocessed SRT task data (high-pass filter of >.009 Hz) and nuisance signals were regressed out (top eight aCompCor components, six motion parameters, and first temporal derivative of motion parameters) using a python-based tool (Tambini & Gorgolewski, 2020). Time-series from the hippocampus (bilateral) were extracted and correlations were computed between the hippocampus and voxels in each participant’s whole-brain mask, with temporal censoring (i.e. motion scrubbing) performed (time points surrounding high-motion events of FD >.5 mm, one TR before through two TRs after (Power et al., 2015)). High-motion runs were excluded from the analysis (seven from sub-2 across four sessions), as determined by runs with mean FD > 2 standard deviations of the mean across sessions (FD cutoff =.22 mm). Correlation maps were Fisher Z-transformed, averaged across scans within a session, and used in a second-level analysis (see <u>ROI definition</u> above).

### Representational similarity analysis approach

Representational similarity analysis (RSA) was used to measure the correlation structure of fractal-evoked activation patterns in each session (Fig. 2A), and assess whether specific hypothesized representations emerged during learning (Fig. 2B) (Kriegeskorte et al., 2008). Activation associated with each unique stimulus was estimated during the stimulus localizer scans. To do so, we constructed separate regressors corresponding to the presentation of each unique stimulus (1.5 s boxcar) using a Least-Squares-All (LSA) approach (Mumford et al., 2014) along with a regressor for repeated images and nuisance regressors (see <u>MRI preprocessing and general analysis methods</u> above). RSA was performed at two levels: across entire ROIs (e.g. hippocampus), or in searchlight cubes within an ROI to measure more localized content. When performing searchlight RSA, all voxels in an ROI served as a potential searchlight center, with the stipulations that: searchlights were bounded by the ROI and whole brain mask, and searchlights containing a small number of voxels were not assessed (those with fewer than ⅓ of total searchlight size). For smaller anatomical (MTL) ROIs, searchlights spanned cubes of 5 × 5 × 5 voxels (functional resolution), with larger searchlight cubes of 7 × 7 × 7 voxels for regions outside of the MTL that spanned larger spatial extents.

For a given ROI or searchlight analysis, multivoxel activation patterns were extracted for each of the 24 stimuli in each participant’s stimulus set, and multivariate noise decomposition was applied (Walther et al., 2016). This uses the residuals time-series from the LSA GLM to account for noise variance in each scan, resulting in activation patterns that are less biased by the noise structure. We then measured the similarity (Fisher Z-transformed correlation) of activation patterns across stimulus pairs (e.g. fractals 1 and 2) between separate fMRI runs or scans (e.g. runs 1 and 4, Fig. 2A). Fisher Z-transformed correlation values were then averaged across all run-pairs for each session, resulting in a correlation or pattern similarity matrix per fMRI session (Fig. 2A right). Particular runs of the stimulus localizer scans were excluded, those with: poor behavioral performance (hit rate on 1-back task < 2 standard deviations from the mean across sessions; one run for sub-1 and two runs for sub-2) or high levels of head motion (eight runs from sub-2, across five sessions), as determined by runs with mean FD > 2 standard deviations of the mean across sessions (FD cutoff =.21 mm). Lastly, outlier Fisher Z-transformed correlation values were excluded prior to averaging correlation values across run-pairs (those exceeding four standard deviations from the mean pattern similarity across all sessions for a given participant).

To measure the presence of structured memory representations, we operationalized two memory representations (Fig. 2B) which were used as predictors of pattern similarity (Fig. 2C). Each hypothesized memory representation was coded using values of +1 or-1 for particular stimulus pairs, and negative values were then re-coded such that the values across all conditions summed to zero (i.e. to enforce equal weighting of positive and negative similarity conditions). These matrices were then used as predictors of pattern similarity values (Fisher Z-transformed correlations), resulting in a beta coefficient estimating ‘pattern strength’ for each fMRI session (Fig. 2C).

Two hypothesized representational structures were examined. A ‘sequence identity’ structure (Fig. 2B left) proposes shared patterns between directly associated fractals (those in the same temporal sequence during the SRT), in contrast to lower similarity between fractals in different temporal sequences. A schematic ‘temporal position’ structure (Fig. 2B right) tests for shared patterns between fractals occupying the same temporal or ordinal sequence position (i.e. first item a sequence) across sequences, as compared to lower between-position similarity. This representation can be thought of as coding for time within a sequence, or reflecting temporal structure that is generalized across sequences.

### Modeling changes in representational strength across learning

To assess changes in pattern similarity across learning, we examined pattern strength (beta estimates) across sessions for a given ROI or searchlight. Pattern strength values for each participant were baseline corrected relative to the first two sessions (before sequence exposure occurred) to specifically examine changes in memory representations over time (Brunec et al., 2020; Schapiro et al., 2012). We used two families of functions to assess reliable changes in the strength of representations over time (shown in Fig. 2D). Logistic or sigmoid shaped functions were used to model changes that developed and remained sustained throughout learning (Fig. 2D, left). Quadratic functions were used to approximate time-limited changes, or those that developed over time and returned to baseline levels present at the beginning of learning (Fig. 2D, right). As shown in Fig. 2D, both families of functions contain a free parameter (the inflection point) that permits flexibility in when in time changes occurred throughout learning. Thus, these sets of functions provided a standardized approach to discovering where in the brain memory representations change over time, while also allowing for characterization of differences in the temporal dynamics across brain regions (i.e. whether changes were captured by logistic or quadratic functions, and comparing the timescales of change via the inflection point). As shown in Fig. 2D, the inflection points of logistic functions capture approximately when in time pattern strength is increasing the most, while inflection points of quadratic functions reflect the peak and initial decline of pattern strength.

Logistic functions took the form: 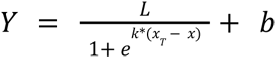 where *Y* is pattern strength and *x* is day in the experiment relative to each participant’s first session (0-81). Four free parameters were fit to the data: the slope (*k*), inflection point (*x_T_*), and baseline (*b*) and asymptote (*L*) values). The free parameters were fit within ranges of:-10 to the maximum × (day) for *x_T_* (permitting values before the first session allows a function with relatively small changes in *Y* that occur early and reach a plateau quickly, initial parameter value = mean x), 0 to 10 for *k* (initial value = 0.1), and 2*min(*Y*) to 2*max(*Y*) for *b* (initial value = 0) and *L* (initial value = mean *Y*-value for 20% of latest data points). Code for the logistic fits is available on github: https://github.com/arielletambini/logistic-model-fitting.

Quadratic functions took the form:

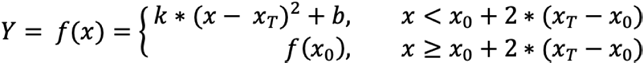

where *x*_0_ is the first day of the experiment, *k* is the 2^nd^ order coefficient, and the other parameters are the same as those described above. A stepwise function was used as it allowed us to model inverted u-shaped changes that occur early in learning without stipulating that pattern strength must continue to decline throughout the remainder of the experiment, instead permitting pattern strength to return to baseline levels. The three free parameters were fit within ranges of: 5 days to 75% of the maximum × (day) for *x_T_* (to ensure that the peak and a portion of the decline in the quadratic function falls within the range of the data, initial value = mean x),-10 to 10 for *k* (initial value = 0.1), and 2*min(*Y*) to 2*max(*Y*) for *b* (initial value = 0).

Scipy (*optimize.curve_fit*) was used to fit all free parameters. For both families of functions, cross-validation was used to assess goodness of model fit and to avoid overfitting. Unlike a typical cross-validation procedure in which each fold of the cross-validation is trained on contiguous data points, here the training and test data for each fold were evenly spaced over time in order to not systematically miss one portion of learning-related changes. Specifically, six cross-validation folds were used and the held out test sessions for each fold were separated by six sessions. For example, in the first cross-validation fold, sessions 1, 7, and 13 were held out while all other sessions were used in model fitting, in the second cross-validation fold, sessions 2, 8, and 14 were held out, and so on. For participants with 18 sessions (sub-2, sub-3), three sessions held out for each of six cross-validation folds. For the participant with 17 sessions (sub-1), three sessions were held out for 5 of 6 folds, and two sessions were held out for one fold. Goodness of model fit was assessed by correlating the held out, predicted levels of pattern strength from each cross-validation fold with observed pattern strength values across all time points. Correlation, as opposed to information-based criterion such as AIC, was used as a metric as it results in an interpretable value of variance explained, enables assessment of statistical significance, and when used in the context of cross-validation, is not biased by the number of free parameters.

Analysis of changes in pattern strength was performed at two levels. When assessing changes in pattern strength at the level of patterns across an entire ROI, i.e. patterns across the left hippocampus (Fig. 3A), we combined pattern strength values across participants in a fixed-effects manner and fit one logistic or quadratic function to all data points (using cross-validation). Significance was assessed via the correlation between predicted and observed pattern strength values across cross-validation folds.

When we performed searchlight analyses (in the native space of each participant), logistic or quadratic functions were fit to the pattern strength values within that searchlight such that they were participant-specific. To assess model fits across searchlights within a given ROI, we computed the sum of the Fisher-Z transformed correlation values across searchlights that showed the effect of interest (e.g. for logistic fits, an increase in pattern strength over time, for quadratic fits, a negative 2^nd^ order coefficient indicating an inverted u-shaped function) divided by the total number of searchlights in the ROI. This value was computed for each participant and then averaged across participants. This metric thus represents a summary measure of goodness-of-fit across searchlights, either for increasing logistic changes over time, or inverted u-shaped quadratic changes. To evaluate the significance of this searchlight metric, permutation testing was used. Null simulations (1000) were performed in which × (day) was shuffled, logistic or quadratic functions were fit to the data in each searchlight, and the same metric described above was computed. This resulted in a null distribution of goodness-of-fit across searchlights for a given ROI, and the true value was compared to this null distribution, resulting in a *p*-value per ROI. False discovery rate (FDR) correction was used to correct for comparisons across multiple ROIs and memory representations (*q* =.05). Specifically FDR correction was applied across all searchlight ROI analyses for a given model type (logistic or quadratic). FDR-corrected *p*-values are reported as *p*_FDR_.

### Timescales of representational changes

After determining which brain regions show reliable changes in memory representations, we examined timescales of pattern similarity changes. As an approximation of when the greatest learning-related change occurred, we examined inflection points of fitted logistic and quadratic functions in searchlights with significant fits, for each ROI that showed at least a trend-level change in pattern strength (*p* < 0.10, FDR-corrected). Specifically, for each ROI, the largest cluster was isolated in each participant that showed an increase in pattern strength over time (cross-validated correlation with *p* < 0.01, minimum of two voxel extent, uncorrected). A less stringent threshold of *p* < 0.05 was used for the angular gyrus and quadratic fits, as these analyses resulted in smaller and relatively few voxels present at *p* < 0.01. Clusters were isolated in all brain regions for two of three participants; one participant (sub-2) showed clusters only for a subset of regions for both sequence (3/5 ROIs) and temporal position (3/6 ROIs) representations. To avoid biasing the results, such that one participant would only contribute to a subset of ROIs, the data from sub-2 were excluded; however, the conclusions were not altered by the inclusion of this data.

Inflection points were estimated for each searchlight using pattern strength values from all sessions to obtain the most accurate estimates (after assessing significance using cross-validation). Inflection points were then pooled across participants in fixed-effects analyses to compare the timescale of changes across ROIs (Fig. 6). To assess whether inflection points differed between brain regions, a nonparametric Kruskall-Wallis one-way analysis of variance (ANOVA) was used. To assess statistical significance, the Kruskall-Wallis test statistic was compared to a null distribution created by shuffling brain region labels and re-calculating the Kruskall-Wallis statistic (100,000 null permutations performed). To test differences in the inflection points between specific pairs of brain regions (e.g. hippocampus vs. neocortical regions), a nonparametric test was used. Specifically, the true difference in mean inflection points across ROIs was computed, and compared to null distributions obtained by permuting brain region labels and re-computing differences in inflection points to assess significance. Finally, to visualize the time course of representational changes across learning, we plotted the idealized model fits across brain regions (Fig. 6). To do so, predicted (idealized) pattern strength values were calculated using the estimated model parameters for each searchlight (using data from all sessions). To facilitate comparisons across brain regions, predicted pattern strength values for logistic fits were normalized from 0 to 1, and averaged across all significant searchlights.

### Informational functional connectivity

To understand whether the expression of distinct memory representations tended to tradeoff on a moment-to-moment basis, we examined trial-by-trial correlations in representational strength across brain regions showing reliable representational changes with learning. To do so, activation was modeled separately for each trial during stimulus localizer scans. Specifically, GLMs contained the following regressors: 48 regressors corresponding to each trial (2 presentations of each stimulus), a regressor for repeated images, and nuisance regressors. We used the same clusters of searchlights as described for the timescale analysis above, focusing on clusters that showed logistic changes over time. In each of these searchlights, we extracted activation patterns across voxels for each trial in the stimulus localizer scans.

To measure pattern strength for each trial (all trials in which a sequence fractal was presented), we first computed the correlation between the pattern for that trial with the average activation patterns for all other stimuli in different runs, yielding a vector of correlations. We then calculated the average correlation across stimulus pairs in positive regions for a given representational matrix (e.g. for sequence identity, the average similarity of a fractal with the other fractals in the same sequence) minus the average correlation across stimulus pairs in negative regions of the representational matrix (e.g. for sequence identity, the average similarity of a fractal with fractals from different sequences). This then yielded a measure of representational strength for each trial in which a sequence stimulus was presented (a total of 24 per run, or 96 per session). This measure was averaged across searchlights within a given ROI to yield one time series of representational strength per ROI and participant. Correlations between trial-by-trial representational strength were then calculated across ROIs within each session, yielding a measure of ‘informational connectivity’ which was then Fisher Z-transformed. We explicitly assessed the strength of representations in ROIs that showed significant development of those representations across learning (e.g. sequence identity representations in posterior fusiform cortex, temporal position representations in parahippocampal cortex).

To ensure that informational connectivity was not driven by other factors that may be shared across trials, such as arousal and attention, we assessed learning-related changes in pattern strength correlations. Specifically, correlations were measured in each session beginning at session 10 (corresponding to ∼36 days, after which inflection points of most logistic functions were observed) and the average correlations in the first two sessions were subtracted (before sequence exposure occurred). To assess the significance of learning-related changes, correlations of a given portion of the correlation matrix (i.e. within sequence identity representations, or correlations across sequence identity representations) were averaged across ROIs within a session and aggregated across sessions and participants in a fixed-effects analysis. One sample t-tests were performed to assess whether learning-related changes were different from zero. Since the goal of this analysis was to assess correlations in memory expression after some extent of representational change occurred, we excluded the entorhinal cortex from this analysis since learning-related changes occurred much later in this region.

## Acknowledgements

We thank Chris Gorgolewski, Regina Lapate, and Ian Ballard for helpful input and discussions and Dan Lurie for assistance with data collection. This work was supported by National Institutes of Health grants MH63901, MH106280, and MH111204.

## Supplementary Figures

**Supplementary Figure 1.**
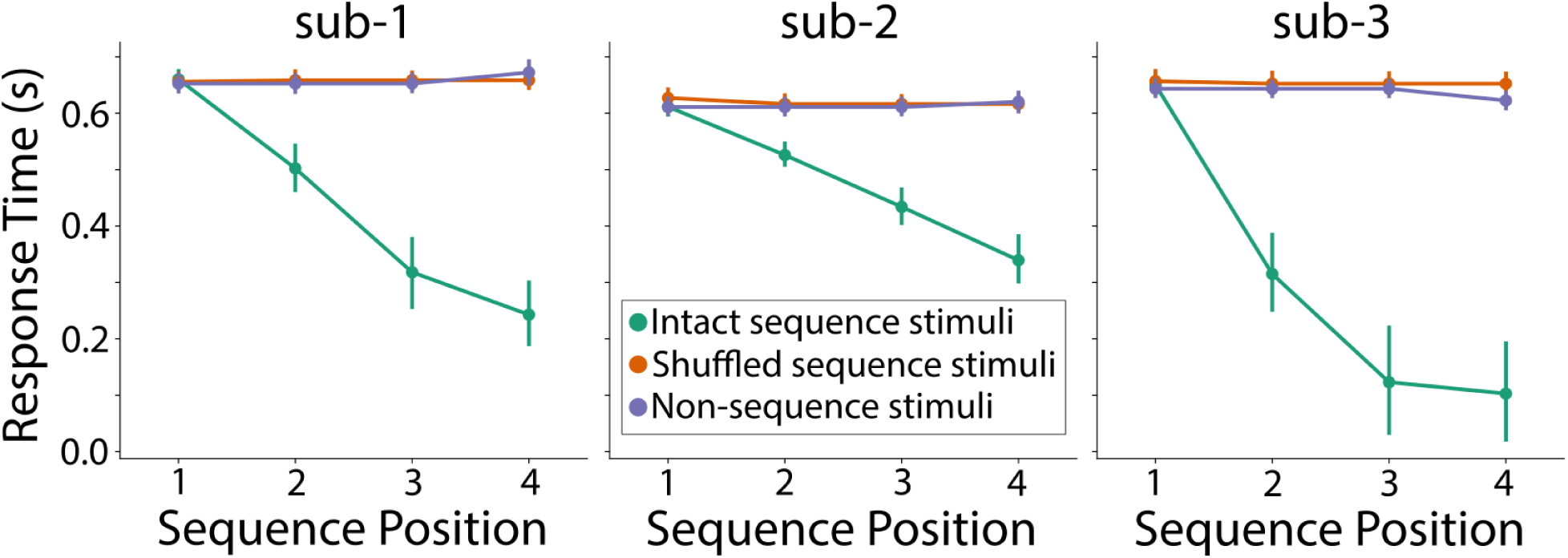
Response times in the SRT task reflect sequence learning. Response time (RT) in the Serial Reaction Time task averaged across all sessions (after sequence exposure began). Responses for sequence stimuli are shown in green (for trials in which intact sequences were shown) and orange (for trials in which sequences were presented in a shuffled order). Non-sequence stimuli are shown in purple. For non-sequence stimuli, the same data are shown for positions 1-3, since positions were not assigned to non-sequence stimuli, with position 4 showing RTs for non-sequence objects (stimulus class which is analogous to sequence stimuli in position 4). Robust reductions in RT are evident for intact sequence stimuli following the first sequence item (position 1).

**Supplementary Figure 2.**
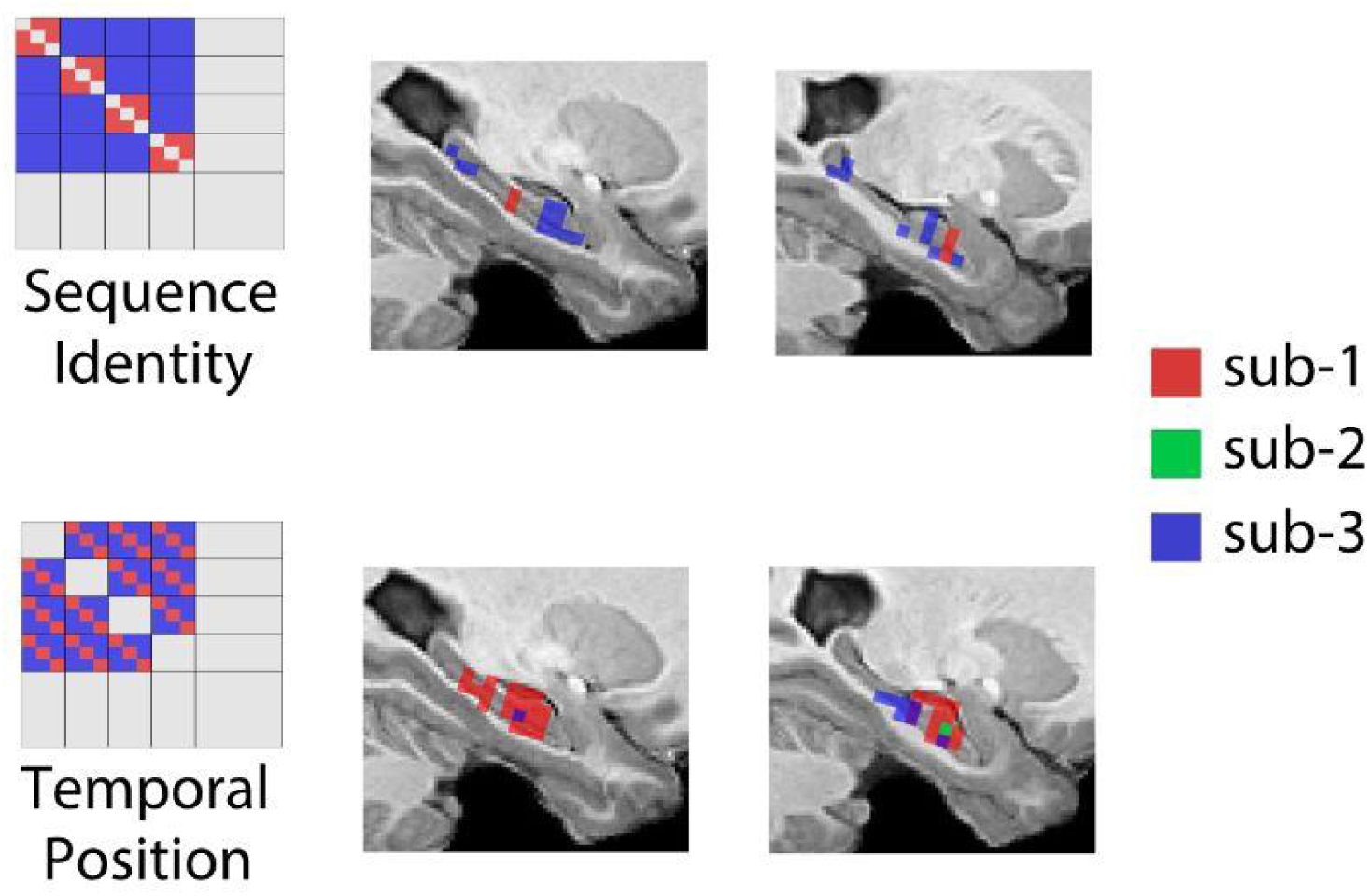
Spatial distribution of hippocampal searchlights showing changes in representational strength with learning. Searchlights in the hippocampus showing changes in representational strength over time (as evidenced by logistic function fits), separately for sequence identity representations (top row) and schematic temporal position representations (bottom row). Voxels indicate searchlight centers that showed significant model fits after False Discovery Rate correction (*q* <.05; correction performed in each participant). Example sagittal slices are shown that contain significant searchlights in multiple participants. Voxels are colored according to each participant (red, sub-1; green, sub-2; blue, sub-3). Data is shown in custom ANTS template space.

**Supplementary Figure 3.**
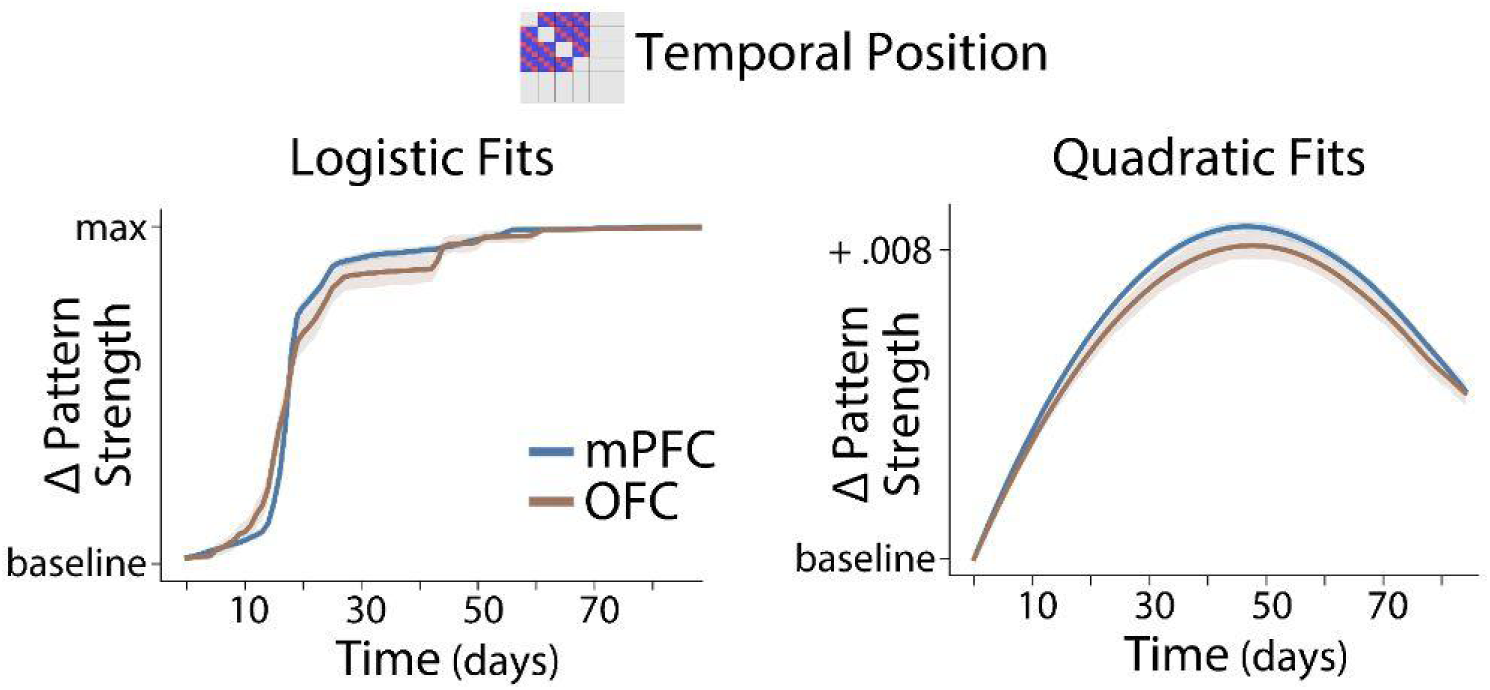
Idealized or best-fit pattern strength across learning in significant searchlights in the mPFC and OFC. Pattern strength values are shown separately for logistic model fits (left plot) and quadratic fits (right plot) for mPFC searchlights (blue) and OFC searchlights (brown). Other conventions are the same as in Figure 6.

## Notes

### Competing Interest Statement

The authors have declared no competing interest.

### Summary of Updates

fixed typos

## References

Abraham, A., Pedregosa, F., Eickenberg, M., Gervais, P., Mueller, A., Kossaifi, J., Gramfort, A., Thirion, B., & Varoquaux, G. (2014). Machine learning for neuroimaging with scikit-learn. Frontiers in Neuroinformatics, 8, 14.

Alvarez, P., & Squire, L. R. (1994). Memory consolidation and the medial temporal lobe: a simple network model. Proceedings of the National Academy of Sciences of the United States of America, 91(15), 7041–7045.

Anagnostaras, S. G., Maren, S., & Fanselow, M. S. (1999). Temporally graded retrograde amnesia of contextual fear after hippocampal damage in rats: within-subjects examination. The Journal of Neuroscience: The Official Journal of the Society for Neuroscience, 19(3), 1106–1114.

Antony, J. W., Ferreira, C. S., Norman, K. A., & Wimber, M. (2017). Retrieval as a fast route to memory consolidation. Trends in Cognitive Sciences, xx, 1–4.

Anzellotti, S., & Coutanche, M. N. (2018). Beyond Functional Connectivity: Investigating Networks of Multivariate Representations. Trends in Cognitive Sciences, 22(3), 258–269.

Audrain, S., & McAndrews, M. P. (2022). Schemas provide a scaffold for neocortical integration of new memories over time. Nature Communications, 13(1), 5795.

Avants, B. B., Tustison, N. J., Song, G., Cook, P. A., Klein, A., & Gee, J. C. (2011). A reproducible evaluation of ANTs similarity metric performance in brain image registration. NeuroImage, 54(3), 2033–2044.

Baldassano, C., Hasson, U., & Norman, K. A. (2018). Representation of Real-World Event Schemas during Narrative Perception. The Journal of Neuroscience: The Official Journal of the Society for Neuroscience, 38(45), 9689–9699.

Batterink, L. J., Paller, K. A., & Reber, P. J. (2019). Understanding the Neural Bases of Implicit and Statistical Learning. Topics in Cognitive Science, 11(3), 482–503.

Bayley, P. J., Hopkins, R. O., & Squire, L. R. (2006). The fate of old memories after medial temporal lobe damage. The Journal of Neuroscience: The Official Journal of the Society for Neuroscience, 26(51), 13311–13317.

Behzadi, Y., Restom, K., Liau, J., & Liu, T. T. (2007). A component based noise correction method (CompCor) for BOLD and perfusion based fMRI. NeuroImage, 37(1), 90–101.

Bellmund, J. L., Deuker, L., & Doeller, C. F. (2019). Mapping sequence structure in the human lateral entorhinal cortex. eLife, 8. https://doi.org/10.7554/eLife.45333

Bellmund, J. L. S., Deuker, L., Montijn, N. D., & Doeller, C. F. (2022). Mnemonic construction and representation of temporal structure in the hippocampal formation. Nature Communications, 13(1), 3395.

Bonnici, H. M., Chadwick, M. J., & Maguire, E. A. (2013). Representations of recent and remote autobiographical memories in hippocampal subfields. Hippocampus, 23(10), 849–854.

Bonnici, H. M., & Maguire, E. A. (2018). Two years later – Revisiting autobiographical memory representations in vmPFC and hippocampus. Neuropsychologia, 110, 159–169.

Bontempi, B., Laurent-Demir, C., Destrade, C., & Jaffard, R. (1999). Time-dependent reorganization of brain circuitry underlying long-term memory storage. Nature, 400(6745), 671–675.

Boorman, E. D., Witkowski, P. P., Zhang, Y., & Park, S. A. (2021). The orbital frontal cortex, task structure, and inference. Behavioral Neuroscience, 135(2), 291–300.

Bornstein, A. M., & Daw, N. D. (2012). Dissociating hippocampal and striatal contributions to sequential prediction learning. The European Journal of Neuroscience, 35(7), 1011–1023.

Brett, M., Markiewicz, C. J., Hanke, M., Côté, M.-A., Cipollini, B., McCarthy, P., Cheng, C. P., Halchenko, Y. O., Cottaar, M., Ghosh, S., Larson, E., Wassermann, D., Gerhard, S., Lee, G. R., Kastman, E., Rokem, A., Madison, C., Morency, F. C., Moloney, B.,…freec. (2020). nipy/nibabel: 2.5.2. https://doi.org/10.5281/zenodo.3745545

Brodt, S., Gais, S., Beck, J., Erb, M., Scheffler, K., & Schönauer, M. (2018). Fast track to the neocortex: A memory engram in the posterior parietal cortex. Science, 362(6418), 1045–1048.

Brodt, S., Pöhlchen, D., Flanagin, V. L., Glasauer, S., Gais, S., & Schönauer, M. (2016). Rapid and independent memory formation in the parietal cortex. Proceedings of the National Academy of Sciences of the United States of America, 113(46), 13251–13256.

Brunec, I. K., Robin, J., Olsen, R. K., Moscovitch, M., & Barense, M. D. (2020). Integration and differentiation of hippocampal memory traces. Neuroscience and Biobehavioral Reviews, 118, 196–208.

Collin, S. H. P., Milivojevic, B., & Doeller, C. F. (2015). Memory hierarchies map onto the hippocampal long axis in humans. Nature Neuroscience, 18(11), 1562–1564.

Coutanche, M. N., & Thompson-Schill, S. L. (2014). Fast mapping rapidly integrates information into existing memory networks. Journal of Experimental Psychology. General, 143(6), 2296–2303.

Cowan, E. T., Liu, A. A., Henin, S., Kothare, S., Devinsky, O., & Davachi, L. (2021). Time-dependent transformations of memory representations differ along the long axis of the hippocampus. Learning & Memory, 28(9), 329–340.

Cox, R. W. (1996). AFNI: software for analysis and visualization of functional magnetic resonance neuroimages. Computers and Biomedical Research, an International Journal, 29(3), 162–173.

Dandolo, L. C., & Schwabe, L. (2018). Time-dependent memory transformation along the hippocampal anterior–posterior axis. Nature Communications, 9(1), 1–11.

Davachi, L. (2006). Item, context and relational episodic encoding in humans. Current Opinion in Neurobiology, 16(6), 693–700.

Deuker, L., Bellmund, J. L., Navarro Schröder, T., & Doeller, C. F. (2016). An event map of memory space in the hippocampus. eLife, 5. https://doi.org/10.7554/eLife.16534

de Voogd, L. D., Fernández, G., & Hermans, E. J. (2016). Awake reactivation of emotional memory traces through hippocampal–neocortical interactions. NeuroImage, 134, 563–572.

Dudai, Y. (2004). The neurobiology of consolidations, or, how stable is the engram? Annual Review of Psychology, 55, 51–86.

Esteban, O., Markiewicz, C. J., Blair, R. W., Moodie, C. A., Isik, A. I., Erramuzpe, A., Kent, J. D., Goncalves, M., DuPre, E., Snyder, M., Oya, H., Ghosh, S. S., Wright, J., Durnez, J., Poldrack, R. A., & Gorgolewski, K. J. (2019). fMRIPrep: a robust preprocessing pipeline for functional MRI. Nature Methods, 16(1), 111–116.

Ezzyat, Y., & Davachi, L. (2014). Similarity breeds proximity: pattern similarity within and across contexts is related to later mnemonic judgments of temporal proximity. Neuron, 81(5), 1179–1189.

Ezzyat, Y., Inhoff, M. C., & Davachi, L. (2018). Differentiation of Human Medial Prefrontal Cortex Activity Underlies Long-Term Resistance to Forgetting in Memory. In The Journal of Neuroscience (Vol. 38, Issue 48, pp. 10244–10254). https://doi.org/10.1523/jneurosci.2290-17.2018

Fernandez, C., Jiang, J., Wang, S.-F., Choi, H. L., & Wagner, A. D. (2023). Representational integration and differentiation in the human hippocampus following goal-directed navigation. eLife, 12. https://doi.org/10.7554/eLife.80281

Frankland, P. W., & Bontempi, B. (2005). The organization of recent and remote memories. Nature Reviews. Neuroscience, 6(2), 119–130.

Fries, P., & Maris, E. (2021). What to do if N is two? https://doi.org/10.48550/arXiv.2106.14562

Furman, O., Mendelsohn, A., & Dudai, Y. (2012). The episodic engram transformed: Time reduces retrieval-related brain activity but correlates it with memory accuracy. Learning & Memory, 19(12), 575–587.

Gais, S., Albouy, G., Boly, M., Dang-Vu, T. T., Darsaud, A., Desseilles, M., Rauchs, G., Schabus, M., Sterpenich, V., Vandewalle, G., Maquet, P., & Peigneux, P. (2007). Sleep transforms the cerebral trace of declarative memories. Proceedings of the National Academy of Sciences of the United States of America, 104(47), 18778–18783.

Ghazizadeh, A., Griggs, W., Leopold, D. A., & Hikosaka, O. (2018). Temporal–prefrontal cortical network for discrimination of valuable objects in long-term memory. In Proceedings of the National Academy of Sciences (Vol. 115, Issue 9). https://doi.org/10.1073/pnas.1707695115

Gilboa, A., & Marlatte, H. (2017). Neurobiology of Schemas and Schema-Mediated Memory. Trends in Cognitive Sciences, 21(8), 618–631.

Gilboa, A., & Moscovitch, M. (2021). No consolidation without representation: Correspondence between neural and psychological representations in recent and remote memory. Neuron, 109(14), 2239–2255.

Gilboa, A., Winocur, G., Grady, C. L., Hevenor, S. J., & Moscovitch, M. (2004). Remembering our past: functional neuroanatomy of recollection of recent and very remote personal events. Cerebral Cortex, 14(11), 1214–1225.

Gilmore, A. W., Quach, A., Kalinowski, S. E., González-Araya, E. I., Gotts, S. J., Schacter, D. L., & Martin, A. (2021). Evidence supporting a time-limited hippocampal role in retrieving autobiographical memories. In Proceedings of the National Academy of Sciences (Vol. 118, Issue 12). https://doi.org/10.1073/pnas.2023069118

Giuliano, A. E., Bonasia, K., Ghosh, V. E., Moscovitch, M., & Gilboa, A. (2021). Differential Influence of Ventromedial Prefrontal Cortex Lesions on Neural Representations of Schema and Semantic Category Knowledge. Journal of Cognitive Neuroscience, 33(9), 1928–1955.

Gordon, E. M., Laumann, T. O., Gilmore, A. W., Newbold, D. J., Greene, D. J., Berg, J. J., Ortega, M., Hoyt-Drazen, C., Gratton, C., Sun, H., Hampton, J. M., Coalson, R. S., Nguyen, A. L., McDermott, K. B., Shimony, J. S., Snyder, A. Z., Schlaggar, B. L., Petersen, S. E., Nelson, S. M., & Dosenbach, N. U. F. (2017). Precision Functional Mapping of Individual Human Brains. Neuron, 95(4), 791–807.e7.

Gorgolewski, K. J., Auer, T., Calhoun, V. D., Craddock, R. C., Das, S., Duff, E. P., Flandin, G., Ghosh, S. S., Glatard, T., Halchenko, Y. O., Handwerker, D. A., Hanke, M., Keator, D., Li, X., Michael, Z., Maumet, C., Nichols, B. N., Nichols, T. E., Pellman, J.,…Poldrack, R. A. (2016). The brain imaging data structure, a format for organizing and describing outputs of neuroimaging experiments. Scientific Data, 3, 160044.

Gorgolewski, K. J., Esteban, O., Markiewicz, C. J., Ziegler, E., Ellis, D. G., Notter, M. P., Jarecka, D., Johnson, H., Burns, C., Manhães-Savio, A., & Others. (2018). Nipype. Software: Practice & Experience.

Goshen, I., Brodsky, M., Prakash, R., Wallace, J., Gradinaru, V., Ramakrishnan, C., & Deisseroth, K. (2011). Dynamics of retrieval strategies for remote memories. Cell, 147(3), 678–689.

Gratton, C., & Braga, R. M. (2021). Editorial overview: Deep imaging of the individual brain: past, practice, and promise. Current Opinion in Behavioral Sciences, 40, iii – vi.

Graves, K. N., Sherman, B. E., Huberdeau, D., Damisah, E., Quraishi, I. H., & Turk-Browne, N. B. (2022). Remembering the pattern: A longitudinal case study on statistical learning in spatial navigation and memory consolidation. Neuropsychologia, 174, 108341.

Greve, D. N., & Fischl, B. (2009). Accurate and robust brain image alignment using boundary-based registration. NeuroImage, 48(1), 63–72.

Harand, C., Bertran, F., La Joie, R., Landeau, B., Mézenge, F., Desgranges, B., Peigneux, P., Eustache, F., & Rauchs, G. (2012). The Hippocampus Remains Activated over the Long Term for the Retrieval of Truly Episodic Memories. In PLoS ONE (Vol. 7, Issue 8, p. e43495). https://doi.org/10.1371/journal.pone.0043495

Haskins, A. L., Yonelinas, A. P., Quamme, J. R., & Ranganath, C. (2008). Perirhinal cortex supports encoding and familiarity-based recognition of novel associations. Neuron, 59(4), 554–560.

Hebscher, M., Wing, E., Ryan, J., & Gilboa, A. (2019). Rapid Cortical Plasticity Supports Long-Term Memory Formation. Trends in Cognitive Sciences, 23(12), 989–1002.

Holdstock, J. S., Mayes, A. R., Isaac, C. L., Gong, Q., & Roberts, N. (2002). Differential involvement of the hippocampus and temporal lobe cortices in rapid and slow learning of new semantic information. Neuropsychologia, 40(7), 748–768.

Huntenburg, J. M., Bazin, P.-L., & Margulies, D. S. (2018). Large-Scale Gradients in Human Cortical Organization. Trends in Cognitive Sciences, 22(1), 21–31.

Jenkinson, M., Bannister, P., Brady, M., & Smith, S. (2002). Improved optimization for the robust and accurate linear registration and motion correction of brain images. NeuroImage, 17(2), 825–841.

Johnson, E. L., Chang, W. K., Dewar, C. D., Sorensen, D., Lin, J. J., Solbakk, A.-K., Endestad, T., Larsson, P. G., Ivanovic, J., Meling, T. R., Scabini, D., & Knight, R. T. (2022). Orbitofrontal cortex governs working memory for temporal order. Current Biology: CB, 32(9), R410–R411.

Kaefer, K., Stella, F., McNaughton, B. L., & Battaglia, F. P. (2022). Replay, the default mode network and the cascaded memory systems model. In Nature Reviews Neuroscience (Vol. 23, Issue 10, pp. 628–640). https://doi.org/10.1038/s41583-022-00620-6

Kark, S. M., & Kensinger, E. A. (2019). Post-Encoding Amygdala-Visuosensory Coupling Is Associated with Negative Memory Bias in Healthy Young Adults. The Journal of Neuroscience: The Official Journal of the Society for Neuroscience, 39(16), 3130–3143.

Kim, H. F., Ghazizadeh, A., & Hikosaka, O. (2015). Dopamine Neurons Encoding Long-Term Memory of Object Value for Habitual Behavior. Cell, 163(5), 1165–1175.

Kim, J. J., & Fanselow, M. S. (1992). Modality-specific retrograde amnesia of fear. Science, 256(5057), 675–677.

Klinzing, J. G., Niethard, N., & Born, J. (2019). Mechanisms of systems memory consolidation during sleep. Nature Neuroscience, 22(10), 1598–1610.

Kragel, P. A., Han, X., Kraynak, T. E., Gianaros, P. J., & Wager, T. D. (2021). Functional MRI Can Be Highly Reliable, but It Depends on What You Measure: A Commentary on Elliott et al. (2020) [Review of Functional MRI Can Be Highly Reliable, but It Depends on What You Measure: A Commentary on Elliott et al. (2020)]. Psychological Science, 32(4), 622–626.

Kriegeskorte, N., Mur, M., & Bandettini, P. (2008). Representational similarity analysis-connecting the branches of systems neuroscience. Frontiers in Systems Neuroscience, 2, 4.

Kumaran, D., Hassabis, D., & McClelland, J. L. (2016). What Learning Systems do Intelligent Agents Need? Complementary Learning Systems Theory Updated. Trends in Cognitive Sciences, 20(7), 512–534.

Laumann, T. O., Gordon, E. M., Adeyemo, B., Snyder, A. Z., Joo, S. J., Chen, M.-Y., Gilmore, A. W., McDermott, K. B., Nelson, S. M., Dosenbach, N. U. F., Schlaggar, B. L., Mumford, J. A., Poldrack, R. A., & Petersen, S. E. (2015). Functional System and Areal Organization of a Highly Sampled Individual Human Brain. Neuron, 87(3), 657–670.

Leshinskaya, A., Nguyen, M., & Ranganath, C. (2022). Integration of event experiences to build relational knowledge in the human brain. In bioRxiv (p. 2022.11.17.516960). https://doi.org/10.1101/2022.11.17.516960

Lewis, C. M., Baldassarre, A., Committeri, G., Romani, G. L., & Corbetta, M. (2009). Learning sculpts the spontaneous activity of the resting human brain. In Proceedings of the National Academy of Sciences (Vol. 106, Issue 41, pp. 17558–17563). https://doi.org/10.1073/pnas.0902455106

Liu, Z.-X., Grady, C., & Moscovitch, M. (2017). Effects of Prior-Knowledge on Brain Activation and Connectivity During Associative Memory Encoding. Cerebral Cortex, 27(3), 1991–2009.

Liu, Z.-X., Grady, C., & Moscovitch, M. (2018). The effect of prior knowledge on post-encoding brain connectivity and its relation to subsequent memory. NeuroImage, 167, 211–223.

MacDonald, C. J., Lepage, K. Q., Eden, U. T., & Eichenbaum, H. (2011). Hippocampal “Time Cells” Bridge the Gap in Memory for Discontiguous Events. Neuron, 71(4), 737–749.

Margulies, D. S., Ghosh, S. S., Goulas, A., Falkiewicz, M., Huntenburg, J. M., Langs, G., Bezgin, G., Eickhoff, S. B., Castellanos, F. X., Petrides, M., Jefferies, E., & Smallwood, J. (2016). Situating the default-mode network along a principal gradient of macroscale cortical organization. Proceedings of the National Academy of Sciences of the United States of America, 113(44), 12574–12579.

Markiewicz, C. J., Gorgolewski, K. J., Feingold, F., Blair, R., Halchenko, Y. O., Miller, E., Hardcastle, N., Wexler, J., Esteban, O., Goncavles, M., Jwa, A., & Poldrack, R. (2021). The OpenNeuro resource for sharing of neuroscience data. eLife, 10. https://doi.org/10.7554/eLife.71774

McClelland, J. L. (2013). Incorporating rapid neocortical learning of new schema-consistent information into complementary learning systems theory. Journal of Experimental Psychology. General, 142(4), 1190–1210.

McClelland, J. L., McNaughton, B. L., & O’Reilly, R. C. (1995). Why there are complementary learning systems in the hippocampus and neocortex: insights from the successes and failures of connectionist models of learning and memory. Psychological Review, 102(3), 419–457.

Miller, J. A., Tambini, A., Kiyonaga, A., & D’Esposito, M. (2022). Long-term learning transforms prefrontal cortex representations during working memory. Neuron. https://doi.org/10.1016/j.neuron.2022.09.019

Mızrak, E., Bouffard, N. R., Libby, L. A., Boorman, E. D., & Ranganath, C. (2021). The hippocampus and orbitofrontal cortex jointly represent task structure during memory-guided decision making. Cell Reports, 37(9), 110065.

Montchal, M. E., Reagh, Z. M., & Yassa, M. A. (2019). Precise temporal memories are supported by the lateral entorhinal cortex in humans. Nature Neuroscience, 22(2), 284–288.

Moore, M., Hu, Y., Woo, S., O’Hearn, D., Iordan, A. D., Dolcos, S., & Dolcos, F. (2014). A comprehensive protocol for manual segmentation of the medial temporal lobe structures. Journal of Visualized Experiments: JoVE, 89. https://doi.org/10.3791/50991

Morton, N. W., Zippi, E. L., Noh, S. M., & Preston, A. R. (2021). Semantic Knowledge of Famous People and Places Is Represented in Hippocampus and Distinct Cortical Networks. The Journal of Neuroscience: The Official Journal of the Society for Neuroscience, 41(12), 2762–2779.

Moscovitch, M., & Gilboa, A. (2021). Systems consolidation, transformation and reorganization: Multiple Trace Theory, Trace Transformation Theory and their Competitors.

Moscovitch, M., Rosenbaum, R. S., Gilboa, A., Addis, D. R., Westmacott, R., Grady, C., McAndrews, M. P., Levine, B., Black, S., Winocur, G., & Nadel, L. (2005). Functional neuroanatomy of remote episodic, semantic and spatial memory: a unified account based on multiple trace theory. Journal of Anatomy, 207(1), 35–66.

Mumford, J. A., Davis, T., & Poldrack, R. A. (2014). The impact of study design on pattern estimation for single-trial multivariate pattern analysis. NeuroImage, 103, 130–138.

Murty, V. P., Tompary, A., Adcock, R. A., & Davachi, L. (2017). Selectivity in Postencoding Connectivity with High-Level Visual Cortex Is Associated with Reward-Motivated Memory. The Journal of Neuroscience: The Official Journal of the Society for Neuroscience, 37(3), 537–545.

Nadel, L., & Moscovitch, M. (1997). Memory consolidation, retrograde amnesia and the hippocampal complex. Current Opinion in Neurobiology, 7(2), 217–227.

Nadel, L., Samsonovich, A., Ryan, L., & Moscovitch, M. (2000). Multiple trace theory of human memory: Computational, neuroimaging, and neuropsychological results. In Hippocampus (Vol. 10, Issue 4, pp. 352–368). https://doi.org/10.1002/1098-1063(2000)10:4<352::aid-hipo2>3.0.co;2-d

Naselaris, T., Allen, E., & Kay, K. (2021). Extensive sampling for complete models of individual brains. Current Opinion in Behavioral Sciences, 40, 45–51.

Naya, Y., & Suzuki, W. A. (2011). Integrating what and when across the primate medial temporal lobe. Science, 333(6043), 773–776.

Naya, Y., Yoshida, M., & Miyashita, Y. (2001). Backward spreading of memory-retrieval signal in the primate temporal cortex. Science, 291(5504), 661–664.

Nee, D. E. (2019). fMRI replicability depends upon sufficient individual-level data. Communications Biology, 2(1), 130.

Newbold, D. J., Laumann, T. O., Hoyt, C. R., Hampton, J. M., Montez, D. F., Raut, R. V., Ortega, M., Mitra, A., Nielsen, A. N., Miller, D. B., Adeyemo, B., Nguyen, A. L., Scheidter, K. M., Tanenbaum, A. B., Van, A. N., Marek, S., Schlaggar, B. L., Carter, A. R., Greene, D. J.,…Dosenbach, N. U. F. (2020). Plasticity and Spontaneous Activity Pulses in Disused Human Brain Circuits. Neuron, 107(3), 580–589.e6.

Patenaude, B., Smith, S. M., Kennedy, D. N., & Jenkinson, M. (2011). A Bayesian model of shape and appearance for subcortical brain segmentation. NeuroImage, 56(3), 907–922.

Popham, S. F., Huth, A. G., Bilenko, N. Y., Deniz, F., Gao, J. S., Nunez-Elizalde, A. O., & Gallant, J. L. (2021). Visual and linguistic semantic representations are aligned at the border of human visual cortex. Nature Neuroscience, 24(11), 1628–1636.

Poppenk, J., Evensmoen, H. R., Moscovitch, M., & Nadel, L. (2013). Long-axis specialization of the human hippocampus. In Trends in Cognitive Sciences (Vol. 17, Issue 5, pp. 230–240). https://doi.org/10.1016/j.tics.2013.03.005

Power, J. D., Mitra, A., Laumann, T. O., Snyder, A. Z., Schlaggar, B. L., & Petersen, S. E. (2014). Methods to detect, characterize, and remove motion artifact in resting state fMRI. NeuroImage, 84, 320–341.

Power, J. D., Schlaggar, B. L., & Petersen, S. E. (2015). Recent progress and outstanding issues in motion correction in resting state fMRI. NeuroImage, 105, 536–551.

Power, J. D., Silver, B. M., Silverman, M. R., Ajodan, E. L., Bos, D. J., & Jones, R. M. (2019). Customized head molds reduce motion during resting state fMRI scans. NeuroImage, 189, 141–149.

Preston, A. R., & Eichenbaum, H. (2013). Interplay of hippocampus and prefrontal cortex in memory. Current Biology: CB, 23(17), R764–R773.

Pruessner, J. C., Li, L. M., Serles, W., Pruessner, M., Collins, D. L., Kabani, N., Lupien, S., & Evans, A. C. (2000). Volumetry of hippocampus and amygdala with high-resolution MRI and three-dimensional analysis software: minimizing the discrepancies between laboratories. Cerebral Cortex, 10(4), 433–442.

Quian Quiroga, R., Kraskov, A., Koch, C., & Fried, I. (2009). Explicit Encoding of Multimodal Percepts by Single Neurons in the Human Brain. Current Biology: CB, 19(15), 1308–1313.

Ralph, M. A. L., Jefferies, E., Patterson, K., & Rogers, T. T. (2017). The neural and computational bases of semantic cognition. Nature Reviews. Neuroscience, 18(1), 42–55.

Ranganath, C., & Ritchey, M. (2012). Two cortical systems for memory-guided behaviour. Nature Reviews. Neuroscience, 13(10), 713–726.

Rempel-Clower, N. L., Zola, S. M., Squire, L. R., & Amaral, D. G. (1996). Three Cases of Enduring Memory Impairment after Bilateral Damage Limited to the Hippocampal Formation. In The Journal of Neuroscience (Vol. 16, Issue 16, pp. 5233–5255). https://doi.org/10.1523/jneurosci.16-16-05233.1996

Richards, B. A., Xia, F., Santoro, A., Husse, J., Woodin, M. A., Josselyn, S. A., & Frankland, P. W. (2014). Patterns across multiple memories are identified over time. Nature Neuroscience, 17(7), 981–986.

Ritchey, M., & Cooper, R. A. (2020). Deconstructing the Posterior Medial Episodic Network. In Trends in Cognitive Sciences (Vol. 24, Issue 6, pp. 451–465). https://doi.org/10.1016/j.tics.2020.03.006

Ritchey, M., Montchal, M. E., Yonelinas, A. P., & Ranganath, C. (2015). Delay-dependent contributions of medial temporal lobe regions to episodic memory retrieval. eLife, 4. https://doi.org/10.7554/eLife.05025

Roy, D. S., Park, Y.-G., Kim, M. E., Zhang, Y., Ogawa, S. K., DiNapoli, N., Gu, X., Cho, J. H., Choi, H., Kamentsky, L., Martin, J., Mosto, O., Aida, T., Chung, K., & Tonegawa, S. (2022). Brain-wide mapping reveals that engrams for a single memory are distributed across multiple brain regions. Nature Communications, 13(1), 1799.

Sakai, K., & Miyashita, Y. (1991). Neural organization for the long-term memory of paired associates. Nature, 354(6349), 152–155.

Sakai, K., Naya, Y., & Miyashita, Y. (1994). Neuronal tuning and associative mechanisms in form representation. In Learning & Memory (Vol. 1, Issue 2, pp. 83–105). https://doi.org/10.1101/lm.1.2.83

Schapiro, A. C., Kustner, L. V., & Turk-Browne, N. B. (2012). Shaping of object representations in the human medial temporal lobe based on temporal regularities. Current Biology: CB, 22(17), 1622–1627.

Schapiro, A. C., Rogers, T. T., Cordova, N. I., Turk-Browne, N. B., & Botvinick, M. M. (2013). Neural representations of events arise from temporal community structure. Nature Neuroscience, 16(4), 486–492.

Schapiro, A. C., Turk-Browne, N. B., Norman, K. A., & Botvinick, M. M. (2016). Statistical learning of temporal community structure in the hippocampus. Hippocampus, 26(1), 3–8.

Schlichting, M. L., Mumford, J. A., & Preston, A. R. (2015). Learning-related representational changes reveal dissociable integration and separation signatures in the hippocampus and prefrontal cortex. Nature Communications, 6, 8151.

Schlichting, M. L., & Preston, A. R. (2014). Memory reactivation during rest supports upcoming learning of related content. Proceedings of the National Academy of Sciences of the United States of America, 111(44), 15845–15850.

Schuck, N. W., Cai, M. B., Wilson, R. C., & Niv, Y. (2016). Human Orbitofrontal Cortex Represents a Cognitive Map of State Space. Neuron, 91(6), 1402–1412.

Schwarzlose, R. F., Swisher, J. D., Dang, S., & Kanwisher, N. (2008). The distribution of category and location information across object-selective regions in human visual cortex. Proceedings of the National Academy of Sciences of the United States of America, 105(11), 4447–4452.

Scoville, W. B., & Milner, B. (1957). Loss of recent memory after bilateral hippocampal lesions. Journal of Neurology, Neurosurgery, and Psychiatry, 20(1), 11–21.

Sekeres, M. J., Winocur, G., & Moscovitch, M. (2018). The hippocampus and related neocortical structures in memory transformation. Neuroscience Letters, 680, 39–53.

Sekeres, M. J., Winocur, G., Moscovitch, M., Anderson, J. A. E., Pishdadian, S., Wojtowicz, J. M., St-Laurent, M., McAndrews, M.-P., & Grady, C. L. (2018). Changes in patterns of neural activity underlie a time-dependent transformation of memory in rats and humans. Hippocampus, 28(10), 303248.

Sherman, B. E., Graves, K. N., & Turk-Browne, N. B. (2020). The prevalence and importance of statistical learning in human cognition and behavior. Current Opinion in Behavioral Sciences, 32, 15–20.

Sinclair, A. H., & Barense, M. D. (2018). Surprise and destabilize: prediction error influences episodic memory reconsolidation. Learning & Memory, 25(8), 369–381.

Sinclair, A. H., & Barense, M. D. (2019). Prediction Error and Memory Reactivation: How Incomplete Reminders Drive Reconsolidation. Trends in Neurosciences, 42(10), 727–739.

Sinclair, A. H., Manalili, G. M., Brunec, I. K., Adcock, R. A., & Barense, M. D. (2021). Prediction errors disrupt hippocampal representations and update episodic memories. Proceedings of the National Academy of Sciences of the United States of America, 118(51). https://doi.org/10.1073/pnas.2117625118

Smith, C. N., & Squire, L. R. (2009). Medial temporal lobe activity during retrieval of semantic memory is related to the age of the memory. The Journal of Neuroscience: The Official Journal of the Society for Neuroscience, 29(4), 930–938.

Sommer, T. (2017). The Emergence of Knowledge and How it Supports the Memory for Novel Related Information. Cerebral Cortex, 27(3), 1906–1921.

Squire, L. R. (1992). Memory and the hippocampus: a synthesis from findings with rats, monkeys, and humans. Psychological Review, 99(2), 195–231.

Squire, L. R., & Alvarez, P. (1995). Retrograde amnesia and memory consolidation: a neurobiological perspective. In Current Opinion in Neurobiology (Vol. 5, Issue 2, pp. 169–177). https://doi.org/10.1016/0959-4388(95)80023-9

Squire, L. R., & Bayley, P. J. (2007). The neuroscience of remote memory. Current Opinion in Neurobiology, 17(2), 185–196.

Squire, L. R., Cohen, N. J., & Nadel, L. (1984). The medial temporal region and memory consolidation: a new hypothesis. In G. Weingartner & E. Parker (Eds.), Memory Consolidation (pp. 185–210). Erlbaum.

Squire, L. R., Genzel, L., Wixted, J. T., & Morris, R. G. (2015). Memory consolidation. Cold Spring Harbor Perspectives in Biology, 7(8), a021766.

Takashima, A., Nieuwenhuis, I. L. C., Jensen, O., Talamini, L. M., Rijpkema, M., & Fernández, G. (2009). Shift from hippocampal to neocortical centered retrieval network with consolidation. The Journal of Neuroscience: The Official Journal of the Society for Neuroscience, 29(32), 10087–10093.

Takashima, A., Petersson, K. M., Rutters, F., Tendolkar, I., Jensen, O., Zwarts, M. J., McNaughton, B. L., & Fernández, G. (2006). Declarative memory consolidation in humans: a prospective functional magnetic resonance imaging study. Proceedings of the National Academy of Sciences of the United States of America, 103(3), 756–761.

Tallman, C. W., Clark, R. E., & Smith, C. N. (2022). Human brain activity and functional connectivity as memories age from one hour to one month. In Cognitive Neuroscience (Vol. 13, Issues 3-4, pp. 115–133). https://doi.org/10.1080/17588928.2021.2021164

Tambini, A., & D’Esposito, M. (2020). Causal Contribution of Awake Post-encoding Processes to Episodic Memory Consolidation. Current Biology: CB, 30(18), 3533–3543.e7.

Tambini, A., & Gorgolewski, K. J. (2020). Denoiser: A nuisance regression tool for fMRI BOLD data. https://doi.org/10.5281/zenodo.4033939

Tambini, A., Ketz, N., & Davachi, L. (2010). Enhanced brain correlations during rest are related to memory for recent experiences. Neuron, 65(2), 280–290.

Thavabalasingam, S., O’Neil, E. B., Tay, J., Nestor, A., & Lee, A. C. H. (2019). Evidence for the incorporation of temporal duration information in human hippocampal long-term memory sequence representations. Proceedings of the National Academy of Sciences of the United States of America, 116(13), 6407–6414.

Thorp, J. N., Gasser, C., Blessing, E., & Davachi, L. (2022). Data-Driven Clustering of Functional Signals Reveals Gradients in Processing Both within the Anterior Hippocampus and across Its Long Axis. In The Journal of Neuroscience (Vol. 42, Issue 39, pp. 7431–7441). https://doi.org/10.1523/jneurosci.0269-22.2022

Tompary, A., & Davachi, L. (2017). Consolidation Promotes the Emergence of Representational Overlap in the Hippocampus and Medial Prefrontal Cortex. Neuron, 96(1), 228–241.e5.

Tompary, A., Zhou, W., & Davachi, L. (2020). Schematic memories develop quickly, but are not expressed unless necessary. Scientific Reports, 10(1), 16968.

Tsao, A., Sugar, J., Lu, L., Wang, C., Knierim, J. J., Moser, M.-B., & Moser, E. I. (2018). Integrating time from experience in the lateral entorhinal cortex. Nature, 561(7721), 57–62.

Tse, D., Langston, R. F., Kakeyama, M., Bethus, I., Spooner, P. a., Wood, E. R., Witter, M. P., & Morris, R. G. M. (2007). Schemas and memory consolidation. Science, 316(5821), 76–82.

Tustison, N. J., Avants, B. B., Cook, P. A., Zheng, Y., Egan, A., Yushkevich, P. A., & Gee, J. C. (2010). N4ITK: improved N3 bias correction. IEEE Transactions on Medical Imaging, 29(6), 1310–1320.

Vanasse, T. J., Boly, M., Allen, E. J., Wu, Y., Naselaris, T., Kay, K., Cirelli, C., & Tononi, G. (2022). Multiple traces and altered signal-to-noise in systems consolidation: Evidence from the 7T fMRI Natural Scenes Dataset. Proceedings of the National Academy of Sciences of the United States of America, 119(44), e2123426119.

van Kesteren, M. T. R., Beul, S. F., Takashima, A., Henson, R. N., Ruiter, D. J., & Fernández, G. (2013). Differential roles for medial prefrontal and medial temporal cortices in schema-dependent encoding: from congruent to incongruent. Neuropsychologia, 51(12), 2352–2359.

van Kesteren, M. T. R., Ruiter, D. J., Fernández, G., & Henson, R. N. (2012). How schema and novelty augment memory formation. Trends in Neurosciences, 35(4), 211–219.

Vezoli, J., Vinck, M., Bosman, C. A., Bastos, A. M., Lewis, C. M., Kennedy, H., & Fries, P. (2021). Brain rhythms define distinct interaction networks with differential dependence on anatomy. Neuron, 109(23), 3862–3878.e5.

Vincent, J. L., Snyder, A. Z., Fox, M. D., Shannon, B. J., Andrews, J. R., Raichle, M. E., & Buckner, R. L. (2006). Coherent spontaneous activity identifies a hippocampal-parietal memory network. Journal of Neurophysiology, 96(6), 3517–3531.

Viskontas, I. V., Carr, V. A., Engel, S. A., & Knowlton, B. J. (2009). The neural correlates of recollection: hippocampal activation declines as episodic memory fades. Hippocampus, 19(3), 265–272.

Visser, R. M., de Haan, M. I. C., Beemsterboer, T., Haver, P., Kindt, M., & Scholte, H. S. (2016). Quantifying learning-dependent changes in the brain: Single-trial multivoxel pattern analysis requires slow event-related fMRI. Psychophysiology, 53(8), 1117–1127.

Walther, A., Nili, H., Ejaz, N., Alink, A., Kriegeskorte, N., & Diedrichsen, J. (2016). Reliability of dissimilarity measures for multi-voxel pattern analysis. NeuroImage, 137, 188–200.

Wammes, J., Norman, K. A., & Turk-Browne, N. (2022). Increasing stimulus similarity drives nonmonotonic representational change in hippocampus. eLife, 11. https://doi.org/10.7554/eLife.68344

Wang, S.-F., Ritchey, M., Libby, L. A., & Ranganath, C. (2016). Functional connectivity based parcellation of the human medial temporal lobe. Neurobiology of Learning and Memory, 134 Pt A(Pt A), 123–134.

Wang, X., Margulies, D. S., Smallwood, J., & Jefferies, E. (2020). A gradient from long-term memory to novel cognition: Transitions through default mode and executive cortex. NeuroImage, 220, 117074.

Waskom, M., Botvinnik, O., Gelbart, M., Ostblom, J., Hobson, P., Lukauskas, S., Gemperline, D. C., Augspurger, T., Halchenko, Y., Warmenhoven, J., Cole, J. B., de Ruiter, J., Vanderplas, J., Hoyer, S., Pye, C., Miles, A., Swain, C., Meyer, K., Martin, M.,…Brunner, T. (2020). mwaskom/seaborn: v0.11.0 (Sepetmber 2020). https://doi.org/10.5281/zenodo.4019146

Wilson, R. C., Takahashi, Y. K., Schoenbaum, G., & Niv, Y. (2014). Orbitofrontal cortex as a cognitive map of task space. Neuron, 81(2), 267–279.

Winocur, G., & Moscovitch, M. (2011). Memory transformation and systems consolidation. Journal of the International Neuropsychological Society: JINS, 17(5), 766–780.

Winocur, G., Moscovitch, M., & Bontempi, B. (2010). Memory formation and long-term retention in humans and animals: convergence towards a transformation account of hippocampal-neocortical interactions. Neuropsychologia, 48(8), 2339–2356.

Yamashita, K.-I., Hirose, S., Kunimatsu, A., Aoki, S., Chikazoe, J., Jimura, K., Masutani, Y., Abe, O., Ohtomo, K., Miyashita, Y., & Konishi, S. (2009). Formation of long-term memory representation in human temporal cortex related to pictorial paired associates. The Journal of Neuroscience: The Official Journal of the Society for Neuroscience, 29(33), 10335–10340.

Yee, E., Chrysikou, E. G., & Thompson-Schill, S. L. (2014). Semantic memory. In K. N. Ochsner (Ed.), The Oxford handbook of cognitive neuroscience, Vol (Vol. 1, pp. 353–374). Oxford University Press, xvi.

Yeo, B. T. T., Thomas Yeo, B. T., Krienen, F. M., Sepulcre, J., Sabuncu, M. R., Lashkari, D., Hollinshead, M., Roffman, J. L., Smoller, J. W., Zöllei, L., Polimeni, J. R., Fischl, B., Liu, H., & Buckner, R. L. (2011). The organization of the human cerebral cortex estimated by intrinsic functional connectivity. In Journal of Neurophysiology (Vol. 106, Issue 3, pp. 1125–1165). https://doi.org/10.1152/jn.00338.2011

Zeithamova, D., de Araujo Sanchez, M.-A., & Adke, A. (2017). Trial timing and pattern-information analyses of fMRI data. NeuroImage, 153, 221–231.

Zeng, T., Tompary, A., Schapiro, A. C., & Thompson-Schill, S. L. (2021). Tracking the relation between gist and item memory over the course of long-term memory consolidation. eLife, 10. https://doi.org/10.7554/eLife.65588

Zhang, Y., Brady, M., & Smith, S. (2001). Segmentation of brain MR images through a hidden Markov random field model and the expectation-maximization algorithm. IEEE Transactions on Medical Imaging, 20(1), 45–57.

Zheng, L., Gao, Z., McAvan, A. S., Isham, E. A., & Ekstrom, A. D. (2021). Partially overlapping spatial environments trigger reinstatement in hippocampus and schema representations in prefrontal cortex. Nature Communications, 12(1), 6231.

Zhou, J., Jia, C., Montesinos-Cartagena, M., Gardner, M. P. H., Zong, W., & Schoenbaum, G. (2021). Evolving schema representations in orbitofrontal ensembles during learning. Nature, 590(7847), 606–611.

Zola-Morgan, S. M., & Squire, L. R. (1990). The primate hippocampal formation: evidence for a time-limited role in memory storage. Science, 250(4978), 288–290.

Zou, F., Guo, W., Allen, E. J., Wu, Y., Charest, I., Naselaris, T., Kay, K., Kuhl, B. A., Hutchinson, J. B., & DuBrow, S. (2022). Re-expression of CA1 and entorhinal activity patterns preserves temporal context memory at long timescales. In bioRxiv. https://doi.org/10.1101/2022.08.31.506090

